# Genetic loci associated with coronary artery disease harbor evidence of selection and antagonistic pleiotropy

**DOI:** 10.1101/064758

**Authors:** Sean G. Byars, Qin Qin Huang, Lesley-Ann Gray, Samuli Ripatti, Gad Abraham, Stephen C. Stearns, Michael Inouye

**Affiliations:** Centre for Systems Genomics, School of BioSciences, The University of Melbourne, Parkville 3010, Victoria, Australia; Department of Pathology, The University of Melbourne, Parkville, Victoria 3010, Australia; National Institute for Health and Welfare, Helsinki, Finland; Wellcome Trust Sanger Institute, Wellcome Trust Genome Campus, Hinxton, Cambridge, United Kingdom; Department of Public Health, University of Helsinki, Helsinki, Finland; Department of Ecology and Evolutionary Biology, Yale University, New Haven, CT, USA

## Abstract

Traditional genome-wide scans for positive selection have mainly uncovered selective sweeps associated with monogenic traits. While selection on quantitative traits is much more common, very few signals have been detected because of their polygenic nature. We searched for positive selection signals underlying coronary artery disease (CAD) in worldwide populations, using novel approaches to quantify relationships between polygenic selection signals and CAD genetic risk. We identified new candidate adaptive loci that appear to have been directly modified by disease pressures given their significant associations with CAD genetic risk. These candidates were all uniquely and consistently associated with many different male and female reproductive traits suggesting selection may have also targeted these because of their direct effects on fitness. This suggests the presence of widespread antagonistic-pleiotropic tradeoffs on CAD loci, which provides a novel explanation for the maintenance and high prevalence of CAD in modern humans. Lastly, we found that positive selection more often targeted CAD gene regulatory variants using HapMap3 lymphoblastoid cell lines, which further highlights the unique biological significance of candidate adaptive loci underlying CAD. Our study provides a novel approach for detecting selection on polygenic traits and evidence that modern human genomes have evolved in response to CAD-induced selection pressures and other early-life traits sharing pleiotropic links with CAD.

**Author Summary:** How genetic variation contributes to disease is complex, especially for those such as coronary artery disease (CAD) that develop over the lifetime of individuals. One of the fundamental questions about CAD — whose progression begins in young adults with arterial plaque accumulation leading to life-threatening outcomes later in life — is why natural selection has not removed or reduced this costly disease. It is the leading cause of death worldwide and has been present in human populations for thousands of years, implying considerable pressures that natural selection should have operated on. Our study provides new evidence that genes underlying CAD have recently been modified by natural selection and that these same genes uniquely and extensively contribute to human reproduction, which suggests that natural selection may have maintained genetic variation contributing to CAD because of its beneficial effects on fitness. This study provides novel evidence that CAD has been maintained in modern humans as a byproduct of the fitness advantages those genes provide early in human lifecycles.

## Introduction

It is well established that modern human traits are a product of past evolutionary forces that have shaped heritable phenotypic and molecular variation, but we are far from understanding what diseases have driven natural selection and how this process has left its imprint across the genome. Although many recent genome-wide multi-population scans have searched for signatures of positive selection [1–9], these studies have detected few signals of selection on candidate loci associated with traits or diseases [10–12]. This suggests that classic ‘selective sweeps’ have been relatively rare in recent human history [13, 14] and that the tools currently used miss most of the smaller selection signals caused by diseases associated with polygenic traits [12]. This limits our understanding of how natural selection has acted on variation underlying complex diseases. In this study, we aimed to comprehensively identify positive selection signals underlying coronary artery disease (CAD) loci with methods designed to detect signals of recent positive selection. We also compared quantitative selection signals in 12 worldwide populations (HapMap3) with patterns of disease risk to identify signals of selection linked to CAD pressure.

Classic population genetics theory describes positive selection with the selective-sweep (or hard-sweep) model, in which a strongly advantageous mutation increases rapidly in frequency (often to fixation) resulting in reduced heterozygosity of nearby neutral polymorphisms due to genetic hitch-hiking [15, 16] and a longer haplotype with higher frequency. Many methods have been developed to detect these signatures [17, 18], including traditional tests that detect differentiation in allele frequencies among population (i.e. Wright’s fixation index, Fst [19]) and more recently developed within population tests for extended haplotype homozygosity (i.e. integrated haplotype score, iHS [9]). Some of the most convincing examples of human adaptive evolution have been uncovered for traits influenced by single loci with large effects. For example, the lactase persistence (*LCT*) and Duffy-null (*DARC*) mutations affecting expression of key proteins in milk digestion [10] and malarial resistance [20] both display hallmarks of selective sweeps. Other loci that are not clearly monogenic but also show selective sweeps are associated with high-altitude tolerance (*EPAS1* [21]) and skin pigmentation (*SLC24A5* and *KITLG* [22]). These previous studies showed that rapid selective sweeps occurred around loci where alleles that were previously rare or absent in populations had large effects on phenotypes.

Motivated by these initial successes and the increasing availability of global population data genotyped on higher resolution arrays (i.e. HapMap Project, 1000 Genomes Project), many genome-wide scans for candidate adaptive loci have recently been performed [11]. These studies suggest that selection may have operated on a variety of biological processes [10] in ways that differ among populations (i.e. local adaptation) [23], has been prevalent in genetic variation linked to metabolic processes [24], and may have often targeted intergenic regions and gene regulatory variants rather than protein-coding regions [12]. However, only the larger signals underlying monogenic traits are typically captured due to the lack of statistical power imposed by the need to correct for genome-wide multiple testing [18]. Most of these candidates also are not yet convincing due to inconsistencies between studies that utilized the same data [14], cannot be validated due to the absence of biological or functional information [25, 26], and perhaps because selective sweeps have actually been rare in human populations [27, 28].

In contrast to population genetics, in quantitative genetics rapid adaptation typically involves selection acting on quantitative traits that are highly polygenic [29, 30]. Under the ‘infinitesimal (polygenic) model’, such traits are likely to respond quickly to changing selective pressures through smaller frequency shifts in many polymorphisms already present in the population [13, 31]. Such alleles would not necessarily sweep to fixation, would produce smaller changes in surrounding heterozygosity, and would thus be hard to detect with most current population genetic methods [14, 26, 32]. Note that polygenic and classic sweep models are not mutually exclusive [13, 33], for alleles with small- and large-effects may both underlie a polygenic trait. Thus the degree to which candidate alleles will be detectable after a selective event will vary. Given that most common diseases are highly polygenic [34], this suggests a need to improve how we detect and understand adaptive signatures in the loci associated with polygenic traits.

Recent selection studies investigating polygenic traits have taken two approaches. The first scans for significant selection signals within genome-wide significant disease effect SNPs. For example, Ding and Kullo [35] found significant population differentiation (Fst) for 8 of 158 index SNPs underlying 36 cardiovascular disease phenotypes, and Raj et al. [36] observed elevated positive selection scores (Fst, iHS) for 37 of 416 index susceptibility SNPs underlying 10 inflammatory-diseases. The second approach tests if aggregated shifts in genome-wide significant allele frequencies are associated with phenotypic differences by population, latitudinal, or environmental gradients, which might indicate local adaptation. For example, Castro and Feldman [37] used 1300 index SNPs underlying many polygenic traits and found elevated adaptive signals (Fst and iHS) above background variation, and Turchin et al. [38] demonstrated moderately higher frequency of 139 height-increasing alleles in a Northern (taller) compared to Southern (shorter) European populations. These approaches all assume that the variants with the most significant p values are the most probable selection targets, but many if not most such variants are tagging tested or untested causal variants, which may themselves be of lower frequencies. This suggests an approach sensitive to more subtle signals of selection and disease risk is needed for polygenic selection.

We chose CAD as a model for examining polygenic selection signals underlying complex disease because it has (and continues to) impose considerable disease burden (selection pressure) in humans [39], its underlying genetic architecture has been extensively studied [40, 41] and many of its risk factors (cholesterol, blood pressure) have been under recent natural selection [42] related to potential pleiotropic effects or tradeoffs with CAD. Antagonistic pleiotropy describes gene effect on multiple linked traits where selection on one may cause fitness tradeoffs (i.e. disease, survival) in the other due to their negative genetic association [43]. Two common misconceptions are that CAD is exclusively late age of onset and only occurs at appreciable frequency in contemporary humans. If that were true, selection might not have had either the opportunity or sufficient time to affect genetic variation associated with CAD. However, CAD manifests early in life [44, 45] and can be detected even in adolescence through degree of atherosclerosis [45, 46] and myocardial infarction events [47]. CAD is also a product of many heritable risk factors (cholesterol, weight, blood pressure) whose variation is expressed during the reproductive period, when CAD could drive selection directly or indirectly. Furthermore, CAD has impacted human populations since at least the ancient Middle Kingdom period, with studies finding the presence of atherosclerosis in Egyptian mummies [48]. This suggests that there has been enough time for genomic signatures of selection related to CAD to develop and be detectable in modern humans.

By combining several 1000 Genomes-imputed datasets including HapMap3 and Finnish SNP data, a large genetic meta-analysis of CAD, and HapMap3 gene expression data, we sought to address the reason(s) why CAD exists in humans by answering the following questions: 1) Has selection recently operated on CAD loci 2) How do selection signals underlying CAD loci vary among populations and are they enriched for gene regulatory effects? 3) Do candidate adaptive signatures overlap directly with CAD genetic risk and is this useful for highlighting disease-linked selection signals? 4) Do CAD-linked selection signals display functional effects and evidence of antagonistic pleiotropy, in that they are also linked to biological processes or traits influencing reproduction?

## Results

To test for selection signals for variants directly linked with CAD, we utilized SNP summary statistics from 56 genome-wide significant CAD loci in Nikpay et al. [41], the most recent and largest CAD case-control GWAS meta-analysis to date, to identify 76 candidate genes for CAD (**Supplementary Materials and Methods**). Nikpay et al. used 60,801 CAD cases and 123,504 controls from a mix of individuals of mainly European (77%), south (13% India and Pakistan) and east (6% China and Korea) Asian, Hispanic and African American (∼4%) descent with genetic variation imputed to a high-density using the 1000 Genomes reference panel. By investigating all SNPs in candidate CAD genes, we aimed to improve detection of smaller polygenic selection signals for the range of functional genic variants and short-range intergenic regulatory variants that would be missed with approaches that only consider genome-wide significant SNPs.

### Signals of positive selection within coronary artery disease loci

We utilised the integrated Haplotype Score (iHS) to estimate positive selection for each SNP underlying candidate CAD genes within each population separately. Because iHS is typically used to detect candidate adaptive SNPs where the selected alleles may not have reached fixation [9], this estimate is well suited for detecting recent signals of selection as opposed to other measures [18]. iHS is also better suited for detecting selection acting on standing variation in polygenic traits [18, 49].

Candidate selection signals were found for many of the 76 CAD genes within each of the 12 worldwide populations (11 HapMap3 populations and Finns; Fig. 1A for top 40 based on their association with CAD log odds genetic risk, Fig. S1 for all 76). These were defined as ‘peaks’ of significantly elevated iHS scores across SNPs within each gene-population combination, with the apex approximating the likely positional target of positive selection.

**Figure 1.**
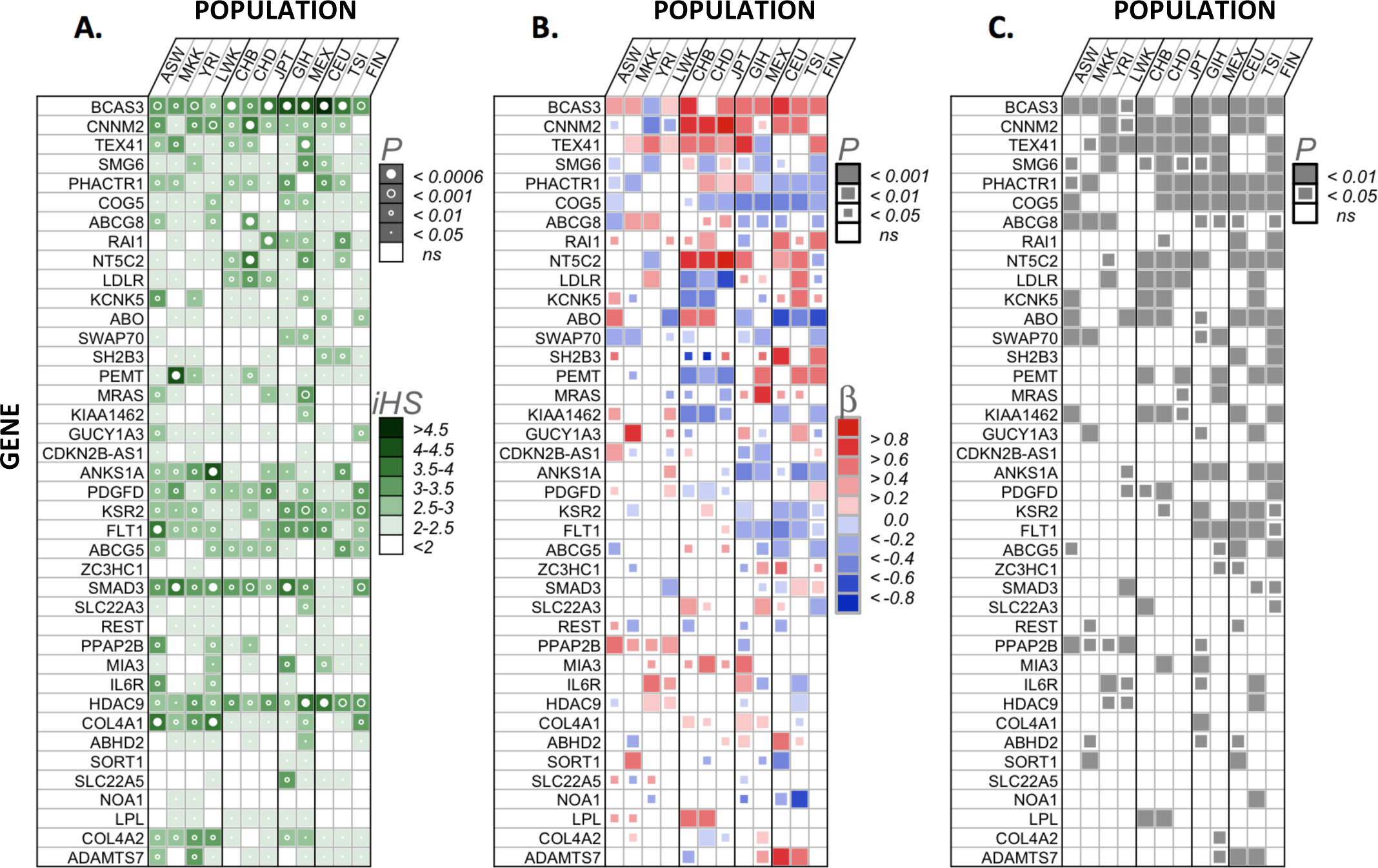
Association of coronary artery disease (CAD) genetic risk and positive signatures of selection in 12 worldwide populations. The 40 of 76 CAD genes investigated are shown that have at least four significant selection-risk associations in Panel B across all 12 populations. **Panel A.** Magnitude and significance of largest positive selection signal (integrated haplotype score, iHS) within each gene-population combination. P values (circles within squares) were obtained from 10000 permutations. Bonferroni corrected p-value limit also shown (α=0.05/76=0.000657) with closed circles. **Panel B.** Null hypothesis: no association between CAD genetic risk and positive selection, tested using mixed effects model with SNP estimates of CAD log odds genetic risk and iHS while accounting for gene LD structure as a random effect (first eigenvector from LD matrix per gene). Scaled regression coefficients were obtained directly from regressions, each p value from 10000 permutations. **Panel C.** Null hypothesis: association between genetic risk and positive selection for SNPs within CAD genes no different than non-CAD associated genes. Permuted p values were estimated by comparing each p value in Panel B against 100 nominal p values obtained by randomly choosing (without replacement) 100 non-CAD associated genes of similar size across the genome and using the same mixed effects model setup as described above. **Populations**. Grouped by ancestry, African (ASW, African ancestry in Southwest USA; MKK, Maasai in Kinyawa, Kenya; YRI, Yoruba from Ibadan, Nigeria; LWK, Luhya in Webuye, Kenya), East-Asian (CHB, Han Chinese subjects from Beijing; CHD, Chinese in Metropolitan Denver, Colorado; JPT, Japanese subjects from Tokyo), European (CEU, Utah residents with ancestry from northern and western Europe from the CEPH collection; TSI, Tuscans in Italy; FIN, Finnish in Finland), GIH (Gujarati Indians in Houston, TX, USA), MEX (Mexican ancestry in Los Angeles, CA, USA).

In the sample of all populations (Fig. 1A, largest iHS scores), most candidate selection signals were relatively small, but a few larger signals were detected. For example, out of the 912 gene-by-population combinations (Fig. S1), 354 (38%) contained weak-moderate candidate selection signals (significant iHS between 2-3), 84 (9%) contained moderate-strong signals (significant iHS between 3-4), and 6 (0.6%) had very strong signals (significant iHS > 4). The 6 largest selection signals were found in the following gene-population combinations: *BCAS3* in GIH (iHS=4.45), MEX (iHS=4.23) and CEU (iHS=4.86), *PEMT* in MKK (iHS=4.24), *ANKS1A* in LWK (iHS=4.03), and *CXCL12* in JPT (iHS=4.10), with all iHS p values <0.0001. Six genes (*BCAS3*, *SMG6*, *PDGFD*, *KSR2*, *SMAD3*, *HDAC9*) exhibited candidate selection signals consistently within all populations (Fig. 1A), and many genes also contained consistent selection signals for all populations within similar ancestral groups (e.g. African, European etc, Fig. 1A).

Within CAD genes, multiple candidate selection signals were sometimes present (particularly within larger genes, within separate linkage disequilibrium (LD)-blocks); these varied between and sometimes within a population. For example, in *PHACTR1* (∼0.57mb in size, 14 introns) there are three main candidate selection signals in introns 4, 7 and 11 (see Fig. S2, comparing cross-population selection signals in *PHACTR1*) that were in separate LD-blocks (see Fig. 3C, LD plots). Within most populations, there was a broad and relatively weak set of candidate selection signals in intron 4 (the largest *PHACTR1* intron, ∼300kb in length). Intron 4 is also the location of the published CAD index SNP (rs9369640) for *PHACTR1*. Three of the African populations had the highest iHS score for the same SNP in intron 4 (rs8180558) including ASW (iHS=2.4, P<0.05), LWK (iHS=2.8, P<0.01) and YRI (iHS=2.2, P<0.05), which is ∼18kb upstream from the index CAD SNP (*r^2^* between rs8180558 and rs9369640 in *PHACTR1*: ASW=0.12; LWK=0.03; YRI=0.04). Peaks of *PHACTR1* selection signals within the three Asian populations were at rs4715043 in CHB (iHS=2.3, P<0.05) and rs6924689 in both CHD (iHS=2.9, P<0.01) and JPT (iHS=3.0,P<0.01). The GIH population contained the largest selection signal, also in intron 4, with an apex at rs4142300 (iHS=3.7, P<0.001, 75kb downstream of/*r^2^*=0.07 with index CAD SNP rs9369640). This corresponded with the same apex SNP in intron 4 for TSI, though the TSI signal was weaker and non-significant (rs4142300, iHS=1.84); rs4142300 was also close to the apex SNP in CEU (rs9349350, iHS=2.0, P<0.05, *r^2^*=0.92) and MEX (rs2015764, iHS=2.1, P<0.05, *r^2^*=0.30). Other significant candidate selection signals were also present in intron 7 for three of the African populations (ASW, LWK, MKK), the CHD and GIH populations, with the largest intron 7 signal within MKK (SNP rs13191209, iHS=3.0, P<0.001). The last significant candidate selection signal within *PHACTR1* was found within intron 11 with the largest signal at rs9349549 (MKK iHS=2.9, P<0.01; CEU iHS=2.7, P<0.01; TSI iHS=3.0, P<0.01). Other interesting candidate selection signals present in other CAD genes (Fig. S1) are not discussed here. Such patterns suggest that candidate selection signals are complex and often do not correspond to the alleles with largest effect on CAD.

### Relationship between CAD genetic risk and selection across populations

For each CAD gene within each population, we used a mixed effects linear model to regress SNP-based estimates of CAD log odds genetic risk (ln(OR), obtained from cardiogramplusc4d.org) against iHS selection scores (**Supplementary Materials and Methods**). We accounted for LD structure by including the first eigenvector from an LD matrix of correlations (*r^2^*) between SNPs within each gene as a random effect.

For a subset of CAD loci, we found significant quantitative associations between disease risk and selection signals and for each of these the direction of this association was often consistent between populations (Fig. 1B). Furthermore, when compared to a null distribution of genes selected randomly from the genome, the strength of the CAD log odds versus selection signal at most loci was statistically significant (Fig. 1C). Fig. 1B shows 40 genes ranked based on those that showed the most consistent number of significant associations across the 12 populations, with those that showed fewer than four significant associations excluded. Positive and negative associations indicate elevated selection signals present in regions with higher or lower CAD log odds genetic risk, respectively.

In the comparison across populations, directionality of significant selection-risk associations tended to be most consistent for populations within the same ancestral group (Fig. 1B). For example, in *PHACTR1*, negative associations were present within all European populations (CEU, TSI, FIN), and in *NT5C2* strong positive associations were present in all East Asian populations (CHB, CHD, JPT). Other negative associations that were consistent across all populations within an ancestry group included five genes in Europeans (*COG5*, *ABO*, *ANKS1A*, *KSR2*, *FLT1*) and four genes (*LDLR*, *PEMT*, *KIAA1462*, *PDGFD*) in East Asians.

Additional consistent positive associations included four genes (*CNNM2*, *TEX41*, *NT5C2*, *MIA3*) in East Asians, three (*BCAS3*, *RAI1*, *KCNK5*) in Europeans, and one (*PPAP2B*) in Africans. In comparison to other ancestral groups, African populations showed fewer significant selection-risk associations (27.9% of all 76-gene x 12-population combinations) than Asians (31.5%) or Europeans (32.8%). Some associations were consistent in all but one population (e.g. *CNNM2*, *ABCG8* in Europeans; *BCAS3*, *KCNK5* in Asians; *CNNM2*, *TEX41* in Africans) or unique to one population within an ancestral group (e.g. *TEX41* in FIN, *COG5* in ASW).

Below we focus on *BCAS3* (Fig. 2) and *PHACTR1* (Fig. 3), two of the strongest selection-risk associations which, when adjusting for LD (**Supplementary Materials and Methods**), displayed varying directionality between at least two populations.

**Figure 2.**
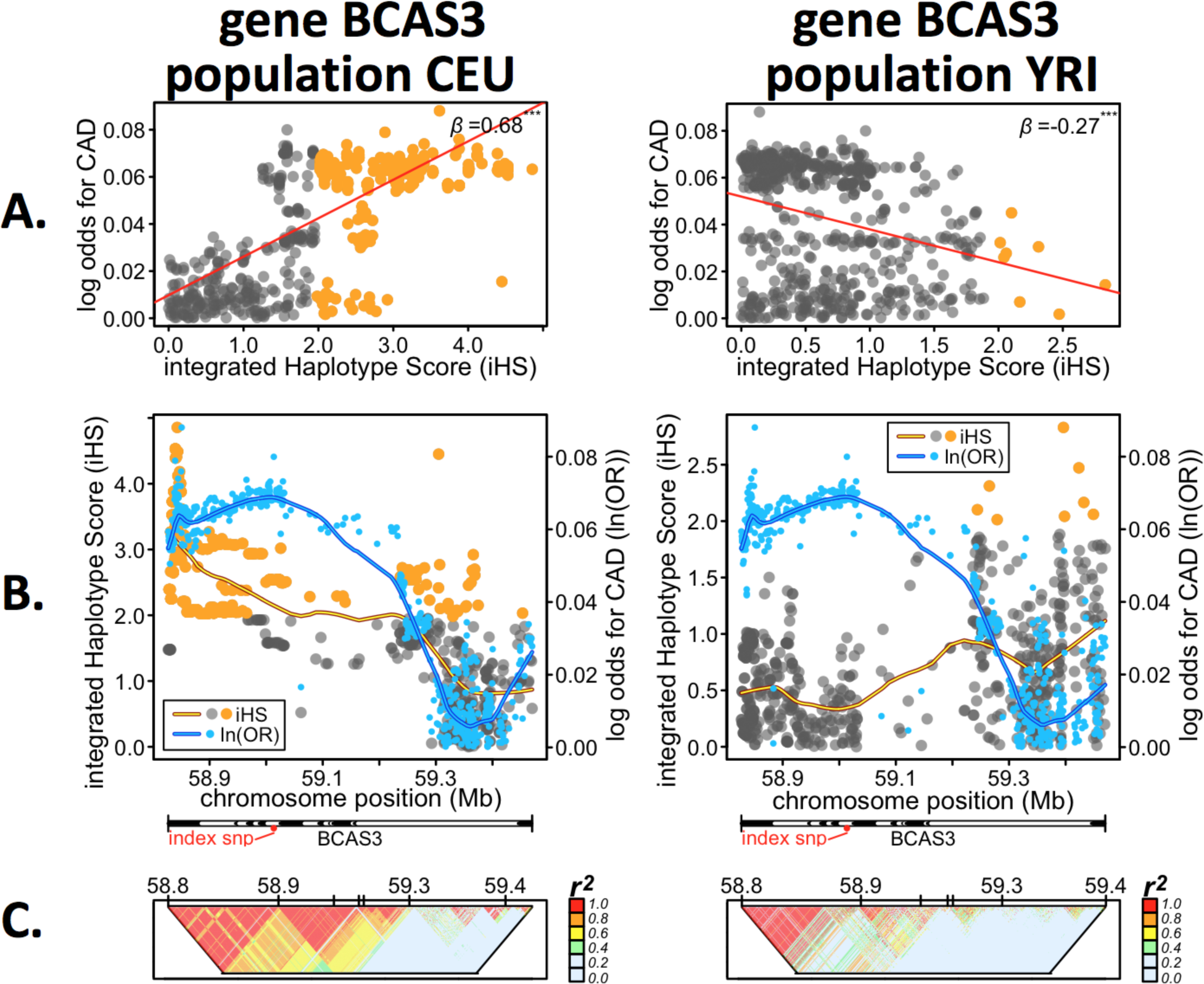
Quantitative links between coronary artery disease risk and selection signals in *BCAS3*. **A.** Correlation between selection signals (iHS) and coronary artery disease (CAD) log odds genetic risk (log odds, ln(OR)), both represented as absolute values. Red line/upper right value, *β* from mixed effects regression. **B.** Base pair positional comparison of selection signals and CAD genetic risk across *BCAS3*. Blue points, CAD log odds values; grey-orange or non-significant-significant points, iHS scores. Horizontal bar shows *BCAS3* gene (and intron) span and location of lead index SNP. Blue/orange lines are smoothed lines estimated with loess function in R. **C.** LD plots, *r^2^*. Populations: CEU, Utah residents with ancestry from northern and western Europe from the CEPH collection; YRI, Yoruba from Ibadan, Nigeria.

**Figure 3.**
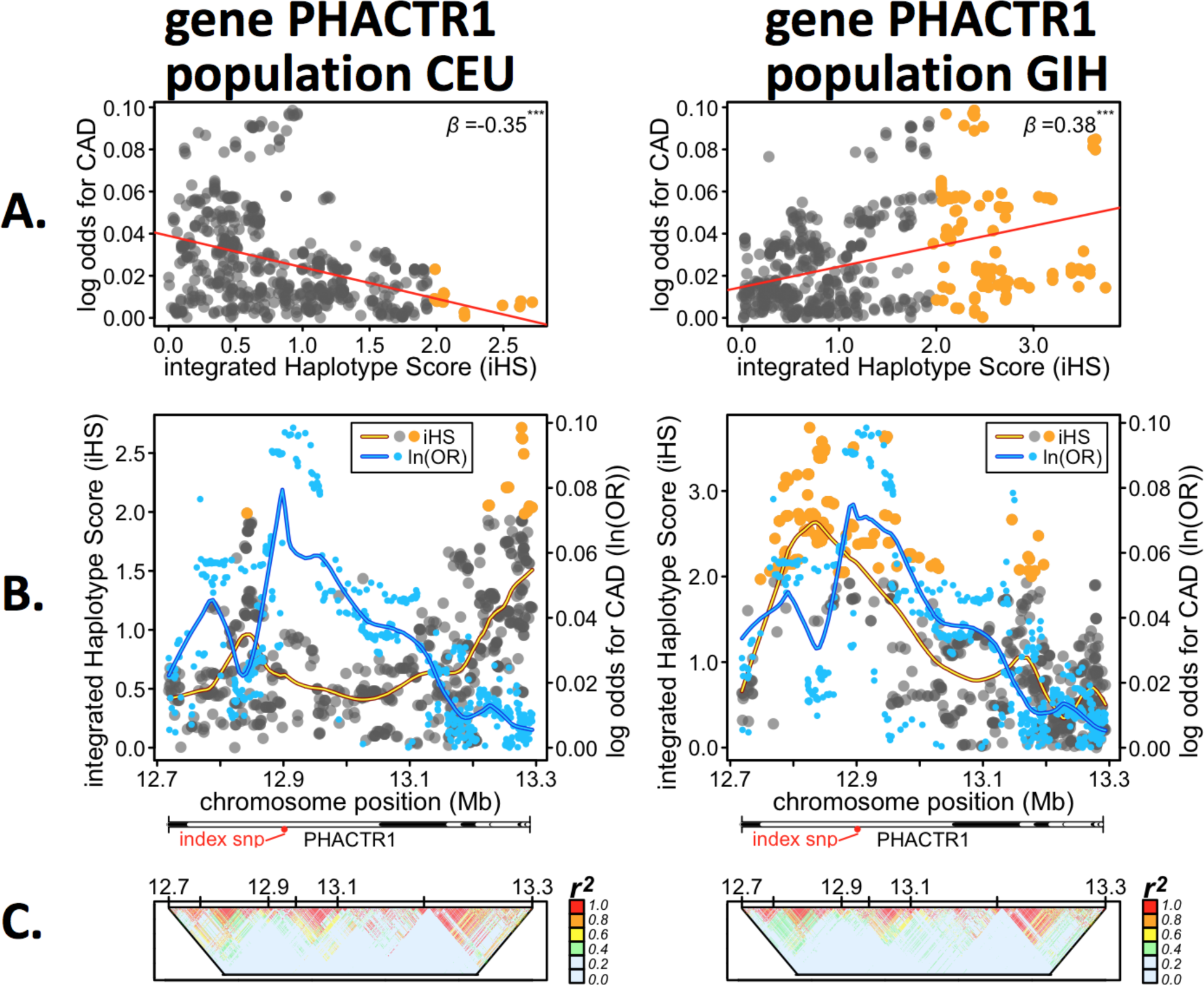
Quantitative links between coronary artery disease risk and selection signals in *PHACTR1*. **A.** Correlation between selection signals (iHS) and coronary artery disease (CAD) log odds genetic risk (ln(OR)), both represented as absolute values. Red line/upper right value, *β* from mixed effects regression. **B.** Base pair positional comparison of selection signals and CAD genetic risk across *PHACTR1*. Blue points, CAD log odds values; grey-orange or non-significant-significant points, iHS scores. Horizontal bar shows *PHACTR1* gene (and intron) spans and location of index SNP if present. **C.** LD plots, *r^2^*. Populations: CEU, Utah residents with ancestry from northern and western Europe from the CEPH collection; GIH, Gujarati Indians in Houston, TX, USA.

#### Genetic risk of CAD vs positive selection in BCAS3

The genetic risk of CAD for variants in *BCAS3* were positively correlated with an extremely large candidate adaptive signal in all European and two of three East Asian populations (Fig. 1B). For example in CEU, the largest iHS score was 4.85 and highly significant, and was elevated across most of *BCAS3* (Fig. 2B CEU, spanning introns 1-18 and various LD-blocks, Fig. 2C), which matched the approximate trends in CAD log odds giving rise to a highly significant positive correlation (Fig. 2A CEU). In contrast, in YRI there was no detectable selection signal close to the index SNP (Fig. 2B YRI), but weak-moderate signals were present towards the end of *BCAS3* (Fig. 2B YRI, introns 18-19, smaller LD-blocks Fig. 2C), which also corresponded with lower CAD log odds (Fig. 2B, YRI) thus giving rise to a significant negative correlation in Fig. 2A.

#### Genetic risk of CAD vs positive selection in PHACTR1

For all European populations, *PHACTR1* (see CEU example, Fig. 3A) selection peaks were typically located within regions of consistently lower CAD log odds (Fig. 3B). This contrasted with most other non-European populations where the highest candidate selection peaks were located within regions with elevated CAD log odds (including the index CAD SNP rs9369640, intron 4). The largest selection peak in GIH (Fig. 3B) overlapped the CAD log odds peak in *PHACTR1* giving rise to the strong positive association seen in Fig. 3A. The two distinctive selection peaks in both CEU and GIH were separated by different LD-blocks (Fig 3C), suggesting that these may have developed independently within *PHACTR1*. Interestingly, the negative association found for the MKK population was due to the location of the selection peaks more closely matching those of the European populations in intron 11 (Fig. S2).

### Enrichment of gene regulatory variants under selection at CAD loci

To establish whether variants with evidence of selection in CAD genes also showed evidence of function, we performed an eQTL scan in 8 HapMap3 populations with matched LCL gene expression. We compared all SNPs in each CAD locus against expression for each focal gene within each population. We found that SNPs with significant integrated Haplotype Scores (iHS) were often also involved in gene regulation, compared to SNPs with non-significant selection scores (Fig. 4, Kolmogorov-Smirnov test p value <0.001). To assess which biological pathways were enriched for the highest-ranked genes according to Fig. 1B, i.e. those where selection scores were most closely associated with CAD log odds genetic risk, we included the top 10 genes into the Enrichr analysis tool [50] and found that these genes are especially enriched in pathways related to metabolism, focal adhesion and transport of glucose and other sugars. More interestingly, we found connections to reproductive phenotypes in the associations of these genes with pathways, ontologies, cell types and transcription factors. For example, we found links to ovarian steroidogenesis and genes expressed in specific cell types and tissues including the ovary, endometrium and uterus (see Table S4 for Enrichr outputs).

**Figure 4:**
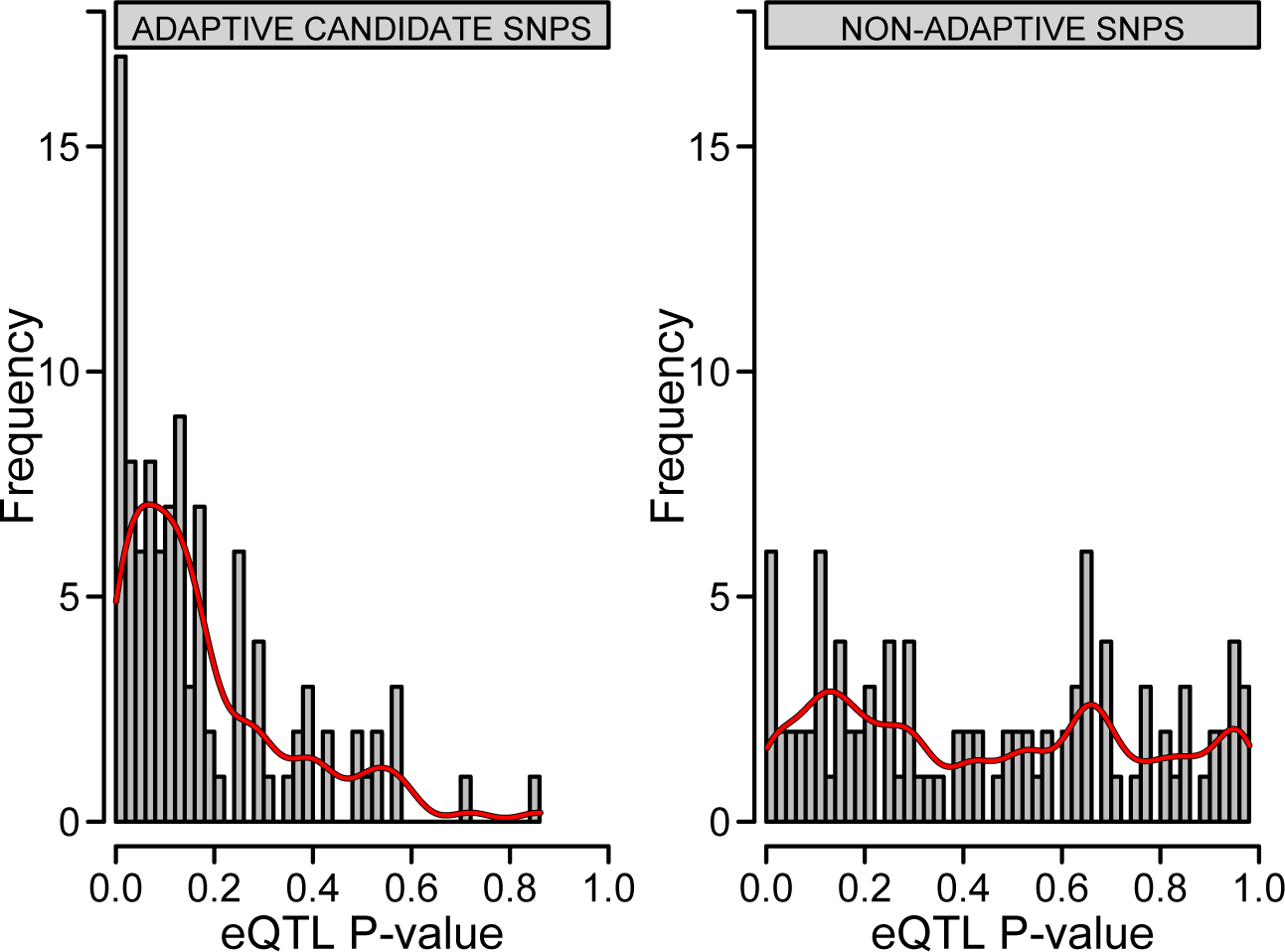
Comparing positive selection with gene regulation. Summary distribution of permuted eQTL p values for SNPs with (left) or without (right) a significant selection signal. SNPs with a significant selection signal (iHS) were chosen by taking the largest significant positive selection signal (if one was present) within each gene-population combination. The same number of SNPs without a significant selection signal were also randomly drawn across all gene-population combinations for comparison. These SNPs were used in an eQTL analysis where they were regressed (including gender as a covariate) against their associated gene probe’s expression.

## Discussion

This study has identified many candidate adaptive signals which suggests that selection on CAD loci is much more widespread than previously appreciated (also see Supplementary Discussion). It has previously been suggested [12] and demonstrated [51] that selection on gene expression levels has been an important element of human adaptation in general. We confirm this result for CAD associated loci. Positive selection signals within CAD loci were more likely than random SNPs to be associated with gene expression levels in *cis* (Fig. 4).

We found evidence that some of these signals may be a result of selection pressures induced directly by CAD itself. This finding is important for highlighting genes that may have been modified directly by selection on disease phenotypes and also for our general understanding of how quickly human genomes can respond to selection induced by changing environments. Subsequent biological process analyses and a thorough literature assessment (below) demonstrated that the loci most consistently associated with CAD genetic risk are also often linked to human reproduction, which suggests both their potential to respond to natural section and their possible role via antagonistic pleiotropy in the reproductive tradeoffs that would help to explain why CAD exists in human populations.

### Coronary artery disease-induced changes to human genomes

One of our most interesting findings was the significant association between selection signals and CAD log odds genetic risk. This approach of integrating genome scans of positive selection with genome-wide genotype-phenotype data has been promoted previously as a tool to uncover biologically meaningful selection signals of recent human adaptation [12, 51] but has rarely been applied. Among the exceptions, Jarvis et al. [52] found a cluster of selection and association signals coinciding on chromosome 3 that included genes *DOCK3* and *CISH*, which are known to affect height in Europeans.

For highly-ranked genes (according to the number of significant associations present within the 12 populations) in Fig. 1B such as *BCAS3*, *CNNM2*, *TEX41*, *SMG6* and *PHACTR1*, the consistent overlap between selection and genetic risk of CAD suggests that many of these may have been modified by CAD-linked selective pressures. If so, then two conditions must have been met. Firstly, CAD was present for long enough to be involved in these genetic alterations, an evolutionary process which generally takes thousands of years. Indeed, precursors of CAD (i.e. atherosclerosis) are detectable in very early civilizations [48]. Secondly, the effects of CAD were directly or indirectly expressed during the reproductive period and trait variation was under natural selection due to its effects on reproductive success.

It is only possible for natural selection to directly act on CAD if those outcomes modify individual fitness relative to others in the same population. As outlined in the introduction, this is possible as CAD outcomes (i.e. myocardial infarction) do occur in young adults. However, early-life CAD outcomes are relatively rare, suggesting selection is more likely to operate indirectly on CAD via its risk factors (or other pleiotropically linked traits, discussed below), which provides a more likely explanation for the close associations we found between positive selection and genetic risk. Supporting this, phenotypic selection has been found operating on CAD risk factors [42], suggesting that these selection pressures are still present in modern humans.

Some genes had large signals of selection but showed weak or no consistent overlap with CAD genetic risk. For example *HDAC9* (Histone Deacetylase 9) shows extensive evidence for having undergone recent selection within most populations, especially those of European or Mexican decent, but little or no overlap with CAD risk was evident in most populations. This suggests positive selection has operated on this gene due to its effects on a trait unrelated to CAD, which may not be surprising given *HDAC9*’s broad biological roles (as a transcriptional regulator, cell-cycle progression) and association with other very different phenotypes including ulcerative colitis [53] and psychiatric disorders [54]. This further demonstrates that this approach is useful for separating candidate selection signals important for the disease or phenotype of interest from those that aren’t.

### Pleiotropic effects that establish the genetic foundations of tradeoffs

To further investigate whether top candidate adaptive loci for CAD modify fitness or share pleiotropic links with other traits that may modify fitness, we performed an extensive systematic literature search on the 40 top-ranked genes in Fig. 1 and a random set of 20 genes. If they have been under selection recently, they might still be associated with reproductive variation (i.e. fitness) in modern environments. We found that all 40 CAD genes shared at least one (often more) connection with fitness (Table S1-S2). Some appear to directly influence fitness (offspring number, age at menarche, menopause, survival), while many were associated with early-life reproductive traits that are likely to indirectly correlate with fitness including variation in ability to fertilize/conceive or fetal growth, development and survival. To test the novelty of this, we randomly chose 20 genes that were approximately the same size as the top 20 genes in Fig. 1. We only found three (out of 20) random genes with at least one potential link with fitness (Table S3). This suggests there are unique pleiotropic links between CAD and traits that have likely been under selection earlier in life.

Evidence for direct links between CAD genes and fitness (Table S1-S2) included genes associated with reproductive (*PPAP2B*, [55]) or twinning (*SMAD3*, [56]) capacity and number of offspring produced (e.g. *KIAA1462*, [57], *SLC22A5*, [58]). *PHACTR1*, *LPL*, *SMAD3*, *ABO* and *SLC22A5* may contribute to reproductive timing (menarche, menopause) in women [59–61] and animals [62]. Expression of *PHACTR1* [63], *KCNK5* [64], *MRAS* and *ADAMST7* [65] appear to regulate lactation capacity. Some gene deficiencies also cause pregnancy loss (e.g. *LDLR*, [66], *COL4A2*, [67]). Evidence for antagonistic links were much more common and included these: 25 genes shared links with traits expressed during pregnancy (Table S1-S2), i.e. variation that can negatively influence the health and survival outcomes of both the fetus and mother [68]. For example, a variant of *CDKN2B-AS1* significantly contributes to risk of fetal growth restriction [69], both *FLT1* [70] and *LPL* [71] are significantly differentially expressed in placental tissues from pregnancies with intrauterine growth restriction (IUGR), and preeclampsia and *LDLR*-deficient mice had litters with significant IUGR [72]. A further 29 and 19 genes were linked to traits that can directly influence female and male fertility, respectively (13 influence both) (Table S1-S2). For example, *BCAS3* and *PHACTR1* are highly expressed during human embryogenesis [73, 74], *SWAP70* is intensely expressed at the site of implantation [75], and *PHACTR1* may play a role in receptivity to implantation [76]. For *ABCG8* and *KSR2,* animal models provide further support as gene expression deficiency can cause infertility in females (*ABCG8*, [77]) and males (*KSR2*, [78]).

Pleiotropic connections were also apparent in the classification of specific disorders or from studies investigating single-gene effects. For example, women with polycystic ovarian syndrome (PCOS) have higher rates of infertility due to ovulation failure and modified cardiovascular disease risk factors (i.e. diabetes, obesity, hypertension [79]). A number of CAD genes in this study (e.g. *PHACTR1*, *LPL*, *PDGFD*, *IL6R*, *CNNM2*) are found differentially expressed in PCOS women [80–84], suggesting possible links between perturbed embryogenesis and angiogenesis. In males, this can be demonstrated with a mutation in *SLC22A5* that causes both cardiomyopathy and male infertility due to altered ability to break down lipids [85, 86]. More generally, many recent studies link altered cholesterol homeostasis with fertility, which is most apparent in patients suffering from hyperlipidemia or metabolic syndrome [87, 88].

To facilitate interpretation of selection occurring on early-life traits or CAD phenotypic risk factors that share pleiotropic connections and possible evolutionary tradeoffs with coronary artery disease, we present a conceptual figure (Fig. 5). These pleiotropic effects are important because many of them affect traits expressed early in life, some extremely early in life. Any allele that increases reproductive performance enough early in life to more than compensate for a loss of associated fitness late in life will be selected [43]. Such a mechanism has been recently suggested to help explain the maintenance of polymorphic disease alleles in modern human populations [89]. Some previous studies have tested for such tradeoffs in humans using direct fitness-related phenotypes (e.g. [90]) although evidence for such a mechanism influencing human disease is currently lacking. Our approach examining antagonistic fitness effects for disease genes that displayed consistent selection-genetic risk associations in diverse worldwide populations provides support for such a mechanism influencing CAD. Here we have presented multiple cases in which such antagonistic pleiotropy appears to be present for genes associated with CAD, which may help to explain our vulnerability to the disease.

**Figure 5.**
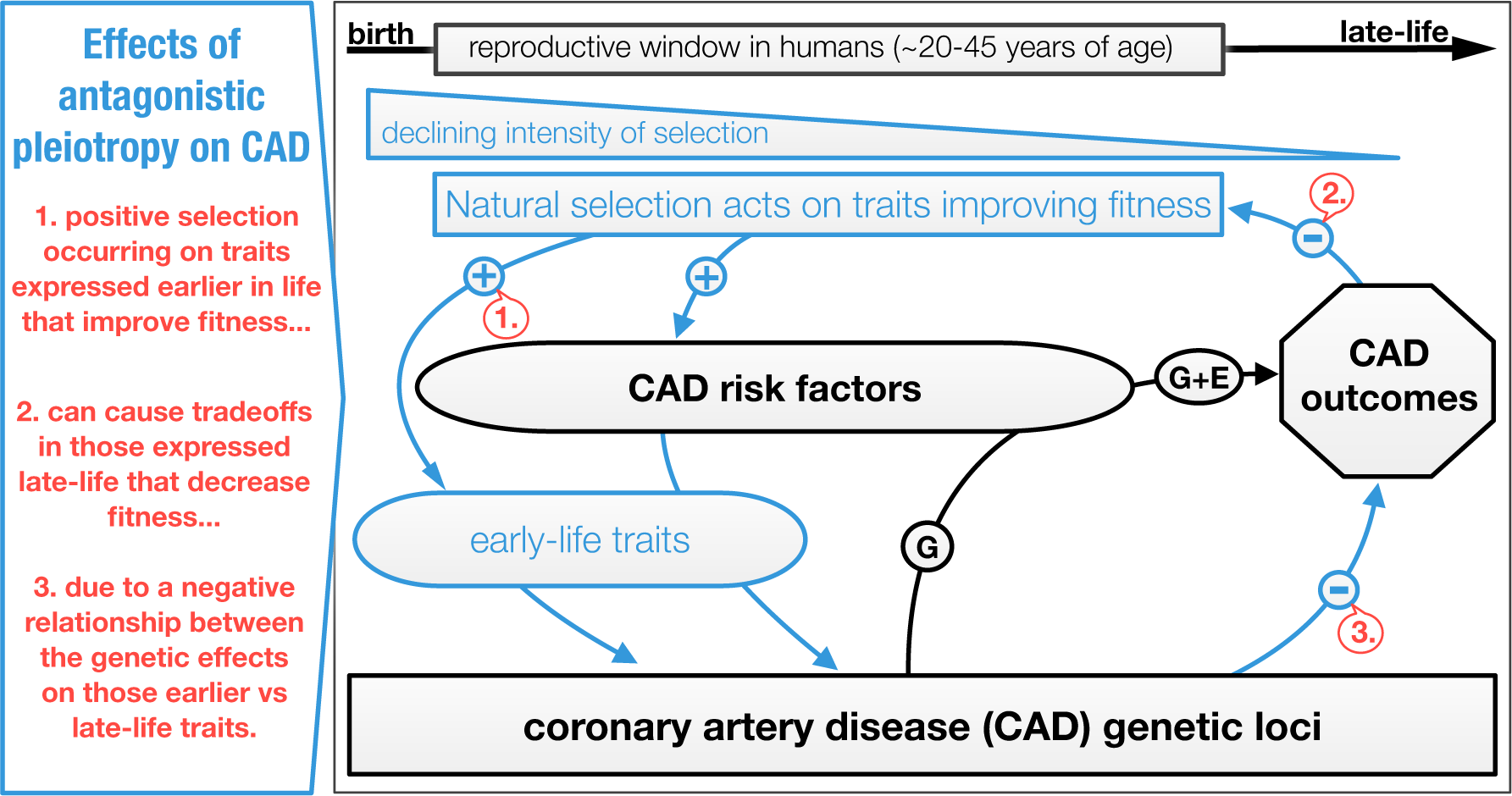
Conceptual figure of potential evolutionary tradeoffs between coronary artery disease (CAD) burden and other phenotypes as a consequence of antagonistic pleiotropy (AP) [43]. As a simple example, AP describes gene effect on two traits (pleiotropy) that oppositely (antagonistic) affect individual fitness at different ages. Selection on that gene conferring a fitness advantage and disadvantage at different ages depends on the size and timing of the effects. An advantage during the ages with the highest probability of reproduction (between∼20-45 years of age in humans) would increase fitness (lifetime reproductive success) more than a similarly sized disadvantage at later ages would decrease it. This concept is part of the well-known evolutionary theory of ageing, which describes tradeoffs in energy invested into growth, reproduction and survival [102]. In the figure above, intense natural selection occurring on CAD loci as a result of fitness advantages (+ signs, red text callout box 1.) conferred by genetically correlated risk factors (‘CAD risk factors’ box) or early-life traits (‘early-life traits’ box) trades off with the deleterious effects of these genes on fitness (i.e. CAD burden) later in life (- sign, red text callout box 2.) where the intensity of selection is weak. This occurs because of the negative relationship between genetic effects on early vs late-life traits (- sign, red text callout box 3.), which could help explain the high prevalence and maintenance of CAD in modern human populations. Over shorter timescales, lifetime probability of CAD is modified by a combination of genetic and environmental risk factors (e.g. [103]). There is a good evidence that such antagonistic effects have operated on CAD loci given: significant associations between CAD genetic risk and selection we found (Fig 1-2); CAD genes also underlie many early-life traits known to modify fitness (Table S2); phenotypic selection has been found operating on CAD phenotypic risk factors [42].

### Study limitations

There are also some limitations to our approach. We utilized CAD genetic risk estimated from a meta-analysis based on predominantly European (77%) with smaller contributions from south/east Asian (19%), Hispanic and African American (∼4%) ancestry [41]. Genetic risk variation for CAD might be different in the un-represented (i.e. Mexican) or less-represented (i.e. African) populations in this meta-analysis. If that were the case, it would reduce the usefulness of comparing selection and risk estimates in those populations. We also saw fewer significant selection-risk associations in the African populations (Fig. 1B), however this may be due to selection signals in the African populations being less obvious than those in East Asian and European populations, perhaps due to lesser linkage disequilibrium, as is consistent with results from previous studies [91]. Calculating disease risk and selection variation from populations within the same ancestral group might help resolve this, however it only represents a potential shortcoming for our cross-population analyses and not observations of antagonistic pleiotropy.

### Summary

In this study, we found evidence that natural selection has recently operated on CAD associated variation. By comparing positive selection variation with genetic risk variation at known loci underlying CAD, we were able to identify and prioritize genes that have been the most likely targets of selection related to this disease across diverse human populations. That selection signals and the direction of selection-risk relationships varied among some populations suggests that CAD-driven selection has operated differently in these populations and thus that these populations might respond differently to similar heart disease prevention strategies. The pleiotropic effects that genes associated with CAD have on traits associated with reproduction that are expressed early in life strongly suggests some of the evolutionary reasons for the existence of human vulnerability to CAD.

## Methods

### Defining loci linked to coronary artery disease

We started with the 56 lead index SNPs from Supplementary Table 5 in Nikpay et al. [41] corresponding to 56 CAD loci. When the index SNP was genic, all SNPs within that gene were extracted (using NCBI’s dbSNP) including directly adjacent intergenic SNPs ±5000bp from untranslated regions (UTR) in LD>0.7 (with any respective genic SNP). When the index SNP was intergenic, that SNP and other directly adjacent SNPs ±5000bp and in LD>0.7 (with the index SNP) were extracted and combined with SNPs from the respective linked gene listed in Nikpay et al. [41] including SNPs ±5000bp from UTR regions in LD>0.7 with that gene. This resulted in SNP lists for 56 genes. To further explore other genes not directly connected with lead index SNPs, but that were found within the CAD loci identified by Nikpay et al. [41], we extracted SNPs within each of those genes (plus SNPs ±5000bp from UTR regions in LD>0.7 with that gene). This resulted in SNP lists for a further 20 genes, bringing the total number of candidate genes for CAD to 76.

The per-SNP log odds (ln(OR)) values for the 76 genes were obtained from Nikpay et al. [41] available at http://www.cardiogramplusc4d.org/downloads and used in the analysis described below.

### Preparation of HapMap3 samples

Genotype data (1,457,897 SNPs, 1,478 individuals) were downloaded for 11 HapMap Phase 3 (release 3) populations (http://www.hapmap.org [92]) including: Yoruba from Ibadan, Nigeria (YRI), Maasai in Kinyawa, Kenya (MKK), Luhya in Webuye, Kenya (LWK), African ancestry in Southwest USA (ASW), Utah residents with ancestry from northern and western Europe from the CEPH collection (CEU), Tuscans in Italy (TSI), Japanese from Tokyo (JPT), Han Chinese from Beijing (CHB), Chinese in Metropolitan Denver, Colorado (CHD), Gujarati Indians in Houston, TX, USA (GIH), and Mexican ancestry in Los Angeles, CA, USA (MEX). We also included another HapMap3 population, the Finnish in Finland (FIN) sample (ftp://ftp.fimm.fi/pub/FIN_HAPMAP3 [93]). These data had already been pre-filtered, i.e. SNPs were excluded that were monomorphic, call rate < 95%, MAF<0.01, Hardy-Weinberg equilibrium P <1x10^-6^ etc.

Before phasing and imputation, we performed a divergent ancestry check with flashpca [94] to check accuracy of population assignments, converted SNP data from build 36 to 37 with UCSC LiftOver (https://genome.ucsc.edu/cgi-bin/hgLiftOver), checked strand alignment in Plink v1.9 [95] to ensure all genotypes were reported on the forward strand, and kept only autosomal SNPs. To speed up imputation, data were first pre-phased with Shapeit v2 [96] using the duoHMM option that combines pedigree information to improve phasing and default values for window size (2Mb), per-SNP conditioning sates (100), effective population size (n=15000) and genetic maps from the 1000 Genomes Phase 3 b37 reference panel (ftp.1000genomes.ebi.ac.uk/vol1/ftp/release/20130502/).

Phased data were imputed in 5 Mb chunks across each chromosome with Impute v2 [97]. We then removed any multiallelic SNPs (insertions, deletions etc) from the imputed data and excluded SNPs with call rate < 95%, HWE P <1x10^-6^ and MAF<1%. The final dataset was then phased with Shapeit v2, and alleles were converted to ancestral and derived states using python script. Ancestral allele states came from 1000 Genomes Project FASTA files and derived 6-primate (human, gorilla, orangutan, chimp, macaque, marmoset) Enredo-Pecan-Ortheus alignment [98] from the Ensembl Compara 59 database [99].

### Estimating signatures of recent selection

*Integrated Haplotype Score (iHS)*: Using the package rehh [100] in R version 3.1.3, per SNP iHS scores were calculated within each population (after excluding non-founders) using methods described previously [9]. iHS could not be calculated for SNPs without an ancestral state, or whose population minor allele frequency is <5%, or for some SNPs that are close to chromosome ends or large regions without SNPs [9]. Rehh was also used to standardize (mean 0, variance 1) iHS values empirically to the distribution of available genome-wide SNPs with similar derived allele frequencies. For analyses in the main text, we considered a SNP to have a candidate selection signal if it had an absolute iHS score > 2, a permuted p value <0.05, and was within a ‘cluster’ of SNPs that also had elevated iHS scores. Although permuting p values is computationally more intensive, it provides more flexibility to detect smaller selection signals that may be incorrectly classified with the more stringent Bonferroni correction that is often applied to these estimates. For the analyses described below, even though we only used iHS estimates for the SNPs defined in the CAD genes (and additional SNPs for permutation purposes), we calculated per-SNP iHS scores genome-wide (rather than locally, i.e. within 1MB regions around focal SNPs), for this provides more accurate estimates because final adjustments are made relative to other genome-wide SNPs of similar sized derived allele frequency classes. P values for iHS scores were permuted based on comparison of nominal p values against 10000 randomly selected estimates from within the same derived allele frequency classes.

### Comparing CAD genetic risk and quantitative selection signals

We first tested the null hypothesis that there is no association between CAD genetic risk and signals of positive selection for CAD genes. For each gene within each population, we used a mixed effects linear model to regress SNP-based estimates of CAD log odds (ln(OR)) genetic risk against selection scores (iHS) resulting in 912 separate regressions. To account for LD structure (and potential confounding of highly correlated SNPs) within each gene, we also included the first eigenvector derived from an LD matrix of correlations (*r^2^*) between SNPs within each gene as a random effect. We chose to model LD structure with mixed-effects models rather than LD-prune because for many genes, the sample would have been too small for regression analyses. Also, it would be very difficult to properly capture both selection and the CAD log odds peaks needed to compare these variables. We accounted for multiple testing by permuting p values for each regression based on comparing each nominal p value against 10000 permuted p values derived from shuffling iHS scores.

Genes were then ranked based on the number of significant associations summed across the 12 populations. The 40 genes with at least four or more significant associations are shown in Fig. 1B. To illustrate the positional architecture of these selection-risk associations, plots for selected highly-ranked genes are shown in Fig. 2-3. By demonstrating how CAD genetic risk peaks and valleys correspond to variation in the magnitude of selection scores (iHS), this allowed visual assessment of potential modifications made to the phenotype-genotype map by selective pressures imposed directly or indirectly by CAD. It also helped us localize selection peaks within genes and compare them between populations. Similar peaks suggested similar selection and different peaks suggested local adaptation. This way of presenting the results also allowed us to detect the smaller adaptive shifts in allele frequencies typically expected to underlie selection on polygenic traits.

We then tested a second null hypothesis: that the selection-risk associations using the CAD genes are not unique compared to non-CAD associated loci. For each of the 76 CAD genes, we randomly (without replacement) chose 100 genes of similar length across the genome and performed the same mixed effects regression procedure described above for each gene by population combination using both CAD log odds values from Nikpay et al. [41], iHS scores estimated from the SNP data, and the first LD eigenvector from SNPs within a gene. Permuted p values were derived by comparing the nominal p value for each CAD gene against the 100 null distribution p values from the non-CAD associated genes. Results are shown in Fig. 1C.

### Identifying functional targets of selection

To examine whether candidate adaptive signals within each gene corresponded to a gene’s regulatory variation, we regressed SNPs within focal genes and gender against that gene’s probe expression levels, which had previously been quantified in lymphoblastoid cell lines using Illumina’s Human-6 v2 Expression BeadChip for eight of the 12 populations [101]. While selection related to CAD may have targeted regulatory variants important for other tissues/cell-types, gene expression data was only available for this cell-type. Given the central importance of circulating lymphoblastoid cells in CAD and its risk factors, we might expect this cell type a good candidate to search for association between selection signals and regulatory variants important for these genes. The raw gene microarray expression data had previously been normalized on a log2 scale using quantile normalization for replicates of a single individual then median normalization for each population [101]. P values for each SNP-probe association were permuted using 10000 permutations by randomly shuffling gene probes expression. P values were then extracted for the most significant iHS score for each gene-population combination and compared to the same number of p values randomly drawn from different LD blocks underlying SNPs with non-significant iHS scores across each gene-population combination. A Kolmogorov-Smirnov test was used to compare the distribution of p values from each. To examine what biological processes were associated with the top ranked genes from Fig. 1, we uploaded the top 10 genes into Enrichr (http://amp.pharm.mssm.edu/Enrichr/) to define associated pathways (i.e. KEGG 2016, kegg.jp/kegg), ontologies (MGI Mammalian phenotypes, informatics.jax.org), cell types (Cancer cell line Encyclopedia, broadinstitute.org/ccle) and transcription factors (ChEA 2015, amp.pharm.mssm.edu/lib/chea.jsp).

## Acknowledgements

This study was supported by the National Health and Medical Research Council (NHMRC) of Australia (grant no. 1062227) and the National Heart Foundation of Australia. MI was supported by a Career Development Fellowship co-funded by the NHMRC and the National Heart Foundation of Australia (no. 1061435). GA was supported by an NHMRC Peter Doherty Early Career Fellowship (no. 1090462). We are grateful to the CARDIoGRAMplusC4D consortium for making their large-scale genetic data available. A list of members of the consortium and the contributing studies is available at www.cardiogramplusc4d.org.

## Author Contributions

Conceptualization, S.G.B. and M.I.; Methodology, S.G.B. and M.I.; Formal analysis, S.G.B. and Q.H.; Literature review, S.G.B.; Writing – original draft, S.G.B. and M.I.; Writing – review & editing, S.G.B., Q.H., L.G., S.R., G.A., S.C.S and M.I.; Visualization, S.G.B.; Funding acquisition, M.I.; Supervision, M.I.

---

**Figure S1:**
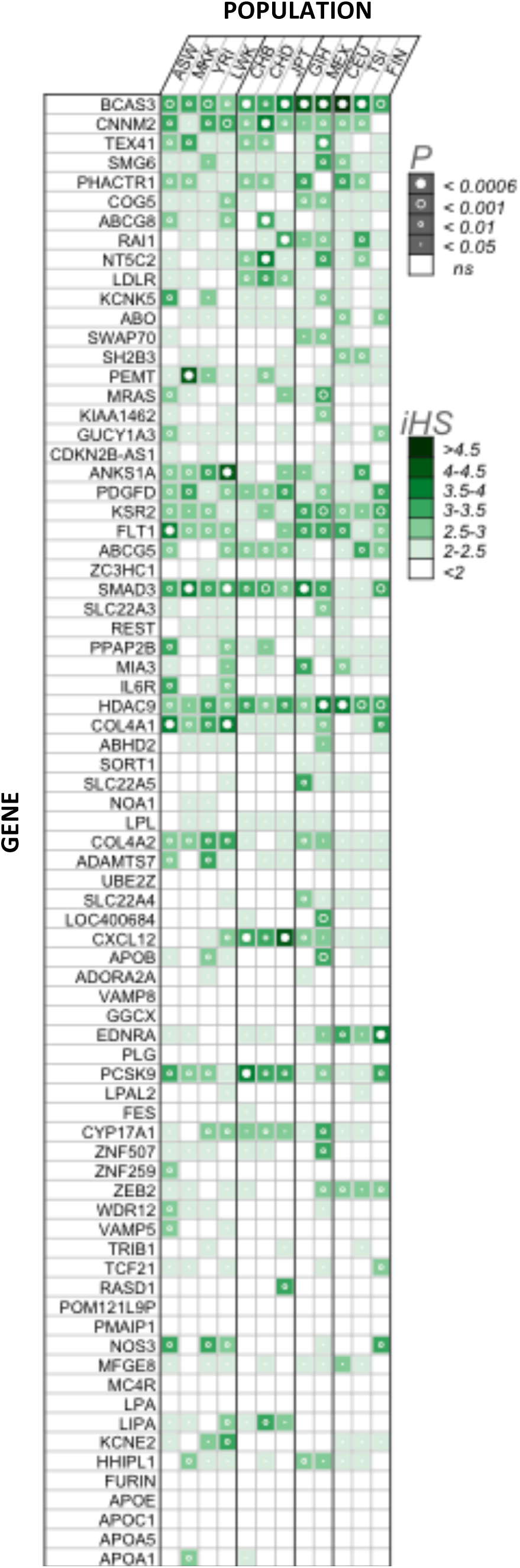
Association of coronary artery disease (CAD) risk and genomic signatures of selection in 12 worldwide populations. All 76 genes are shown ranked according to Fig. 1B. Boxes show magnitude and significance of largest positive selection signal (integrated haplotype score, iHS) within each gene-population combination. P values (circles within squares) were obtained from 10000 permutations. Bonferroni corrected p value limit also shown (α=0.05/76=0.000657) with closed circles. **Populations**. Grouped by common ancestry, African (ASW, African ancestry in Southwest USA; MKK, Maasai in Kinyawa, Kenya; YRI, Yoruba from Ibadan, Nigeria; LWK, Luhya in Webuye, Kenya), East-Asian (CHB, Han Chinese subjects from Beijing; CHD, Chinese in Metropolitan Denver, Colorado; JPT, Japanese subjects from Tokyo), European (CEU, Utah residents with ancestry from northern and western Europe from the CEPH collection; TSI, Tuscans in Italy; FIN, Finnish in Finland), GIH (Gujarati Indians in Houston, TX, USA), MEX (Mexican ancestry in Los Angeles, CA, USA).

**Figure S2:**
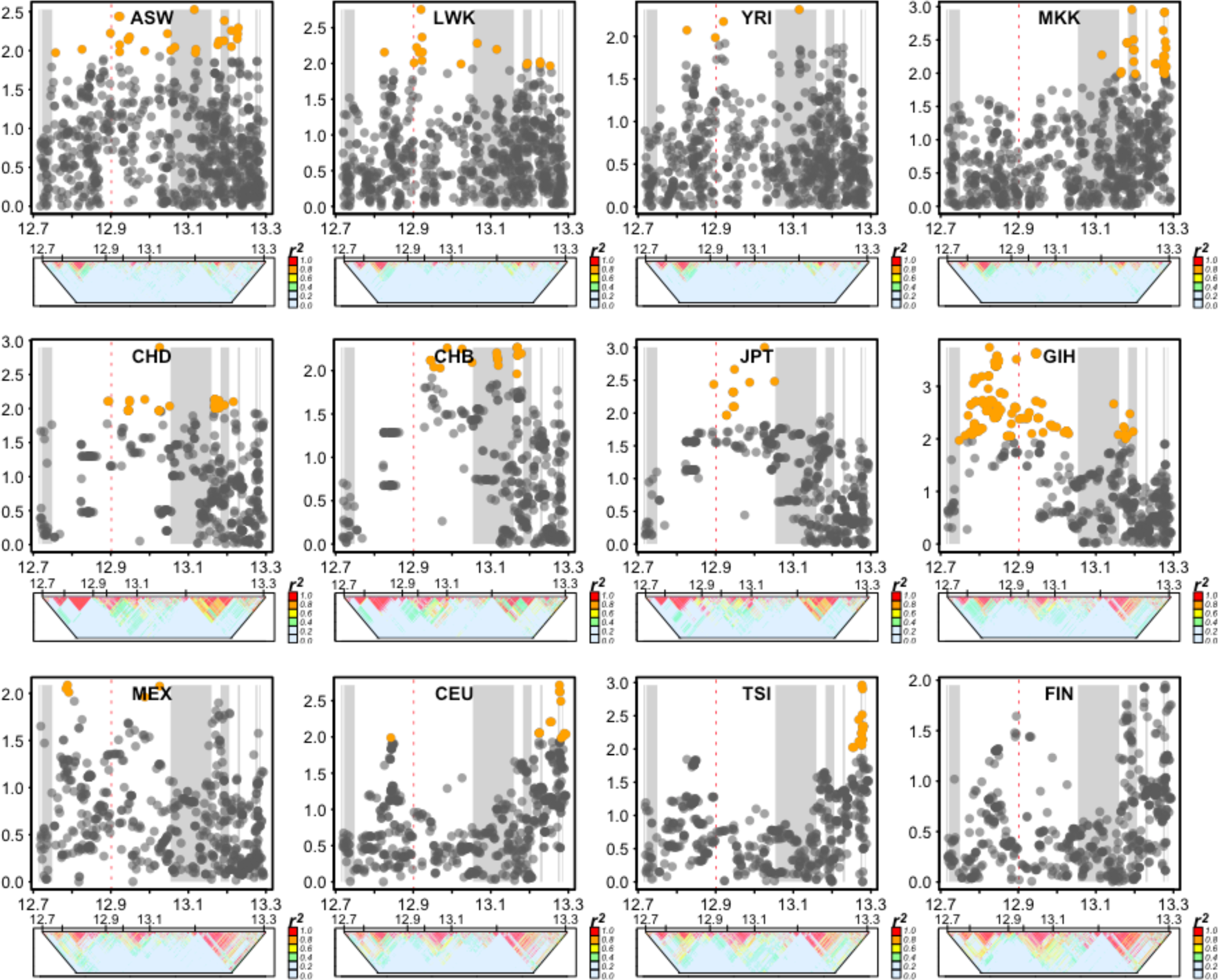
Comparing cross-population candidate selection signals in *PHACTR1*. Per-SNP integrated Haplotype Scores (iHS) plotted by chromosome position within *PHACTR1* (including LD plots below each) for 12 worldwide populations. Permuted p value significance for each score coded by color (grey, non-significant; orange, p < 0.05). Red dashed line indicates position of index SNP for *PHACTR1*. Grey columns in background represent intron spans. Populations are clustered by common ancestry, African (ASW, African ancestry in Southwest USA; MKK, Maasai in Kinyawa, Kenya; YRI, Yoruba from Ibadan, Nigeria; LWK, Luhya in Webuye, Kenya), East-Asian (CHB, Han Chinese subjects from Beijing; CHD, Chinese in Metropolitan Denver, Colorado; JPT, Japanese subjects from Tokyo), European (CEU, Utah residents with ancestry from northern and western Europe from the CEPH collection; TSI, Tuscans in Italy; FIN, Finnish in Finland), GIH (Gujarati Indians in Houston, TX, USA), MEX (Mexican ancestry in Los Angeles, CA, USA).

**Table S1.**
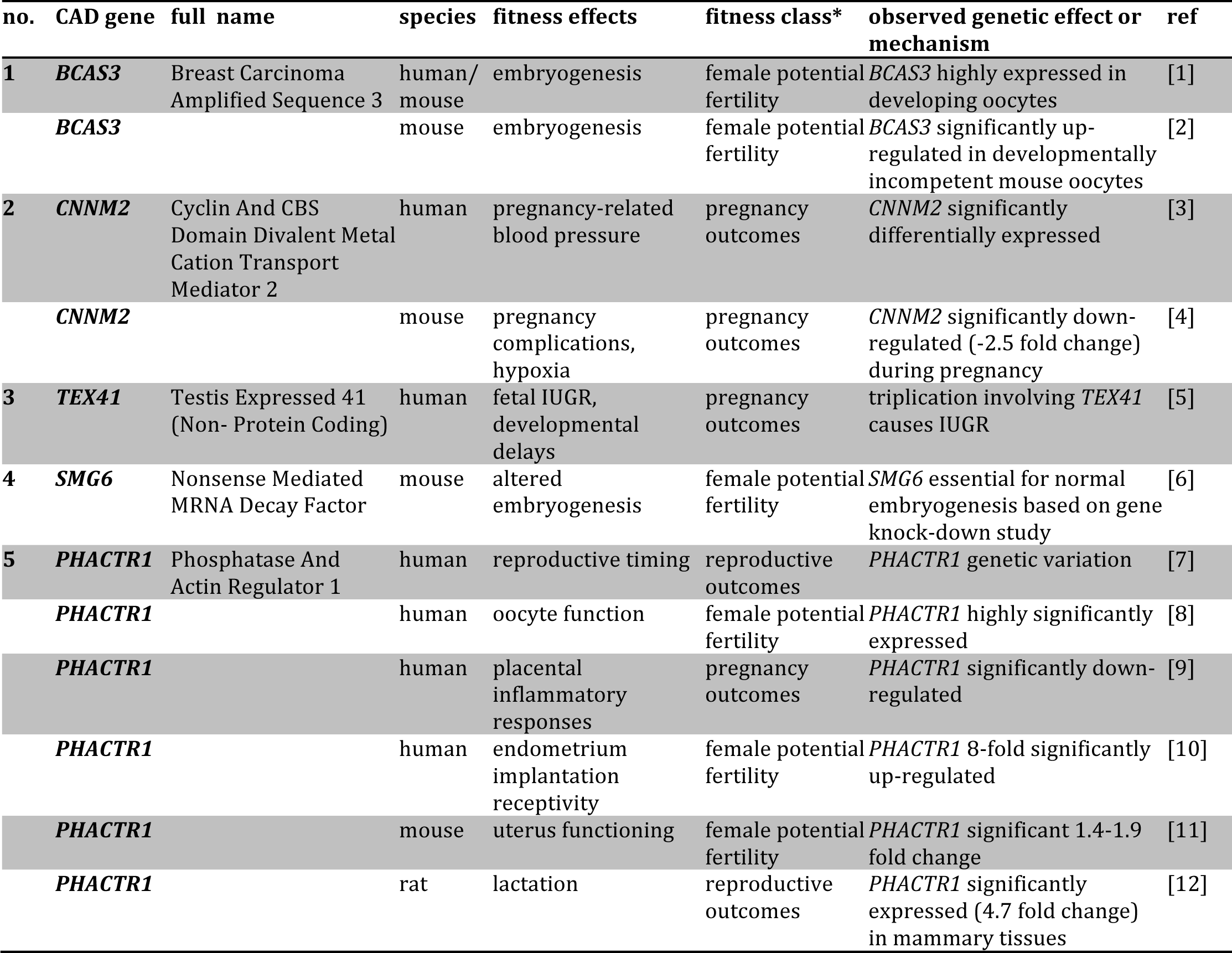
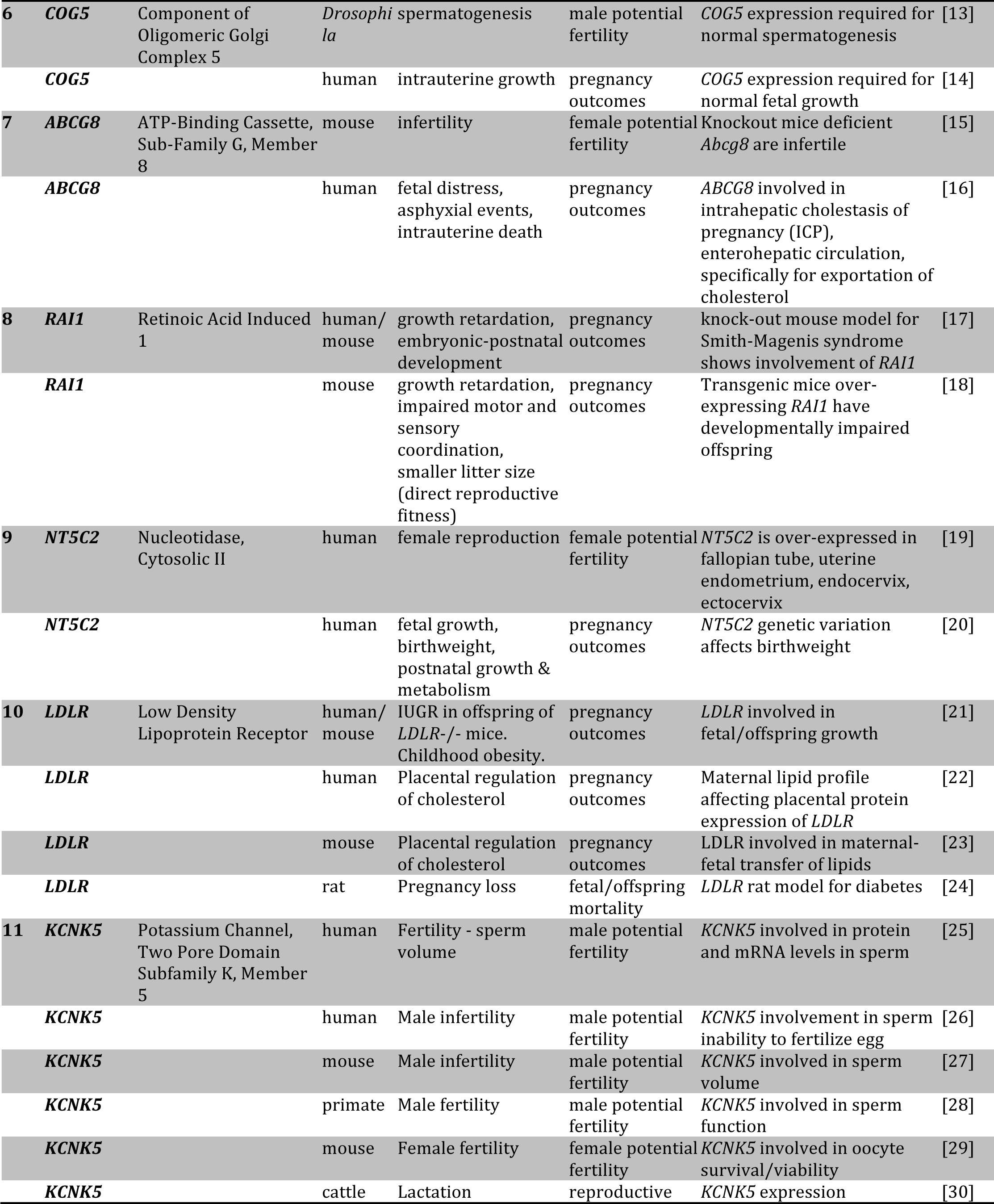
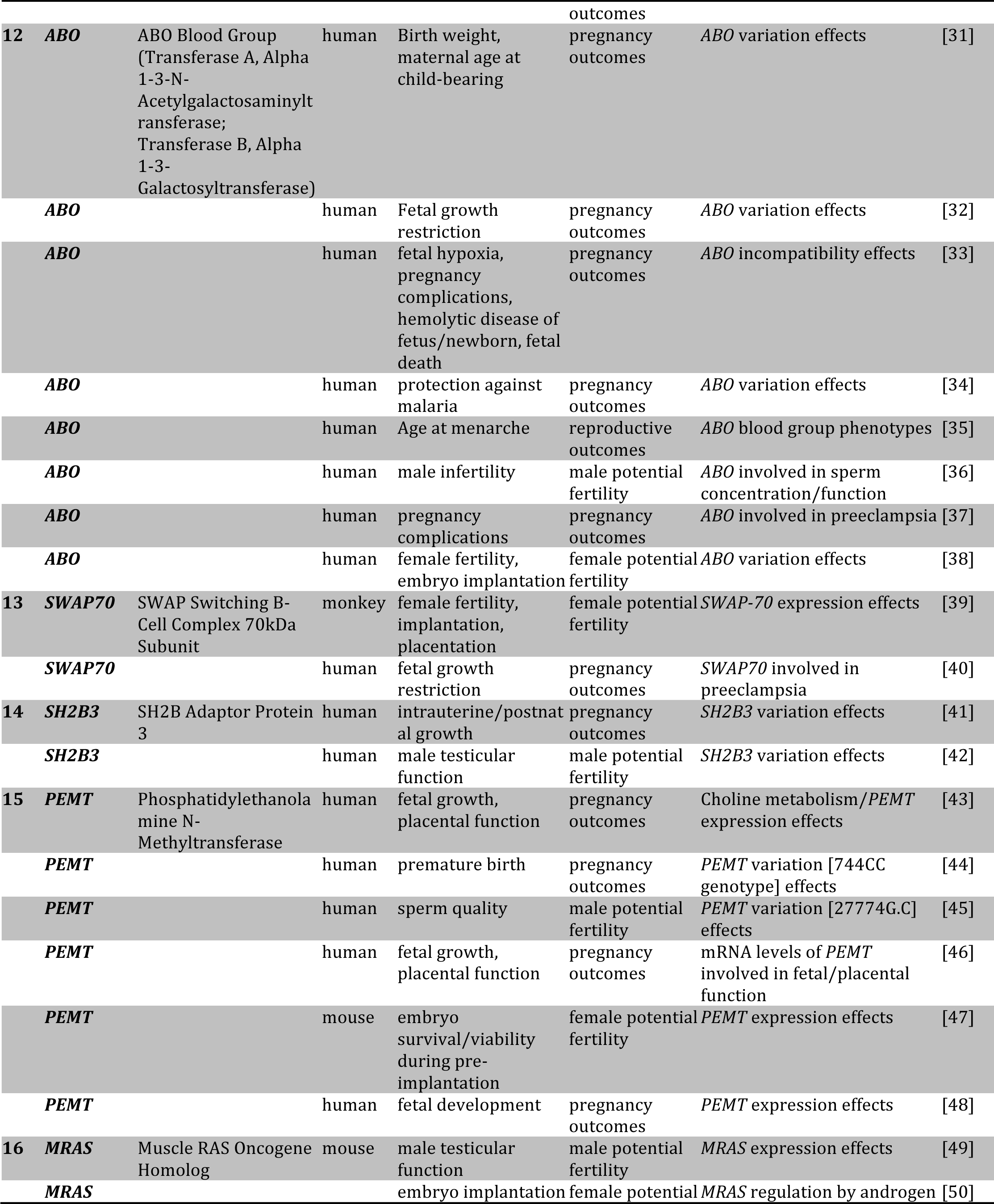
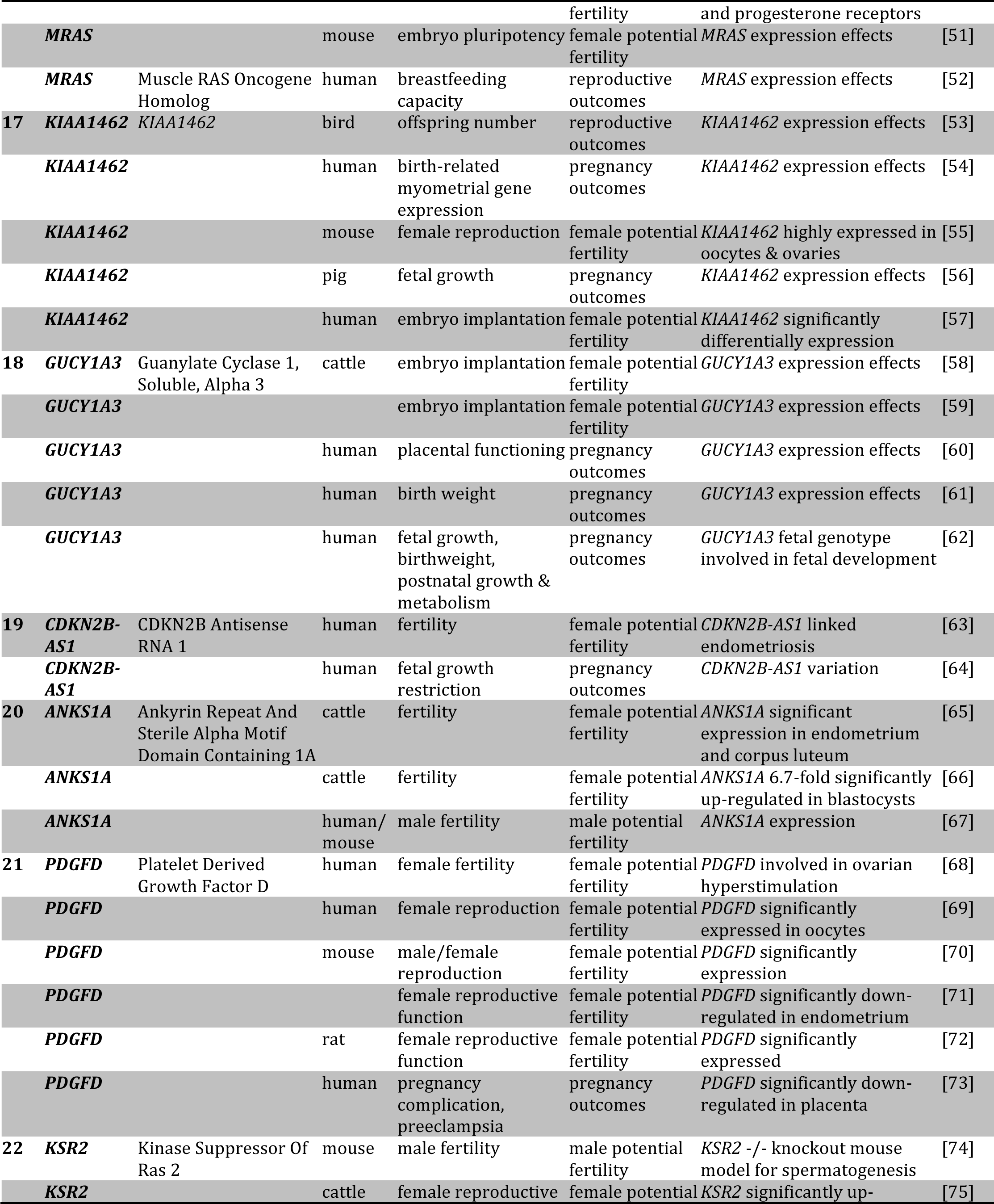
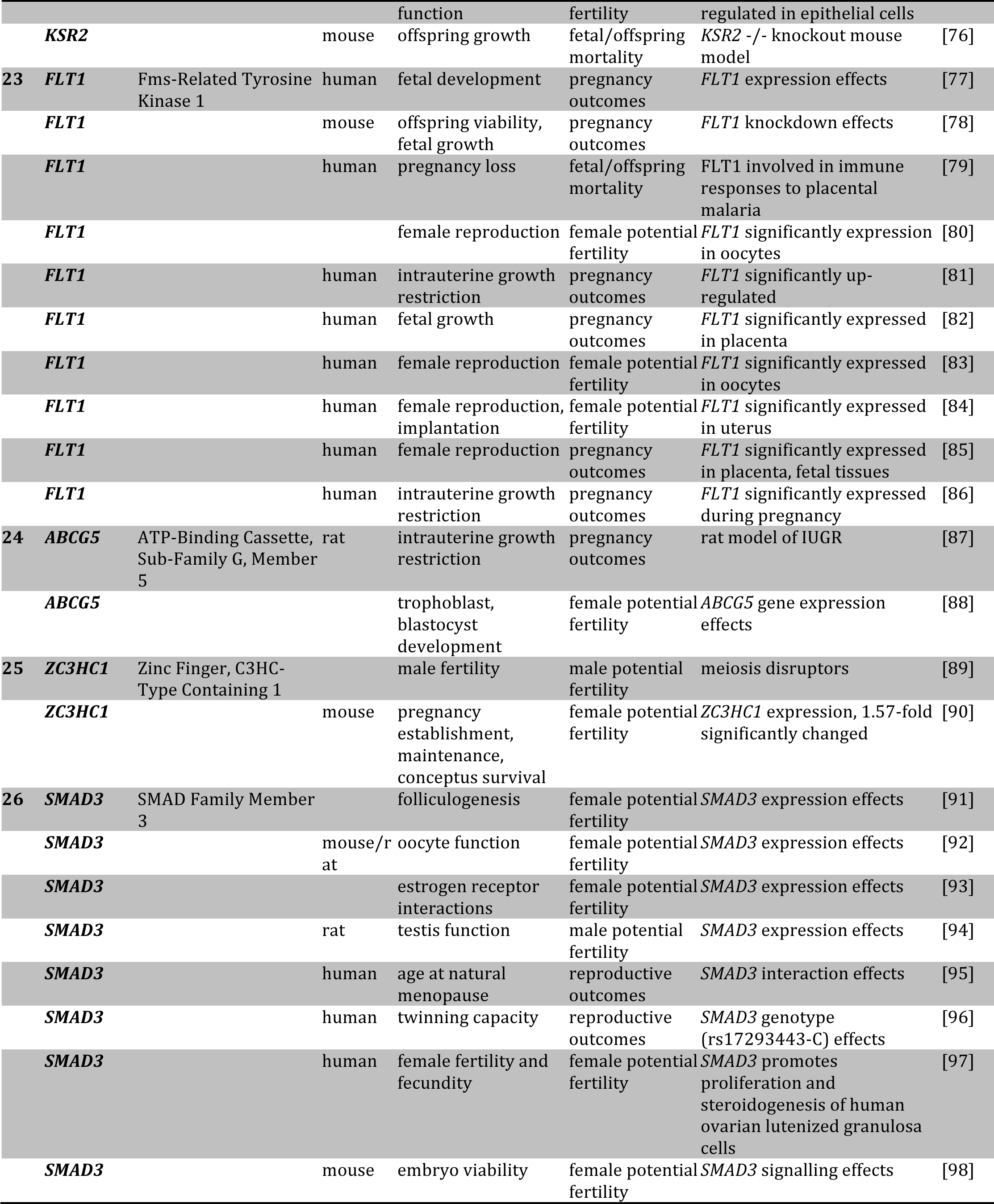
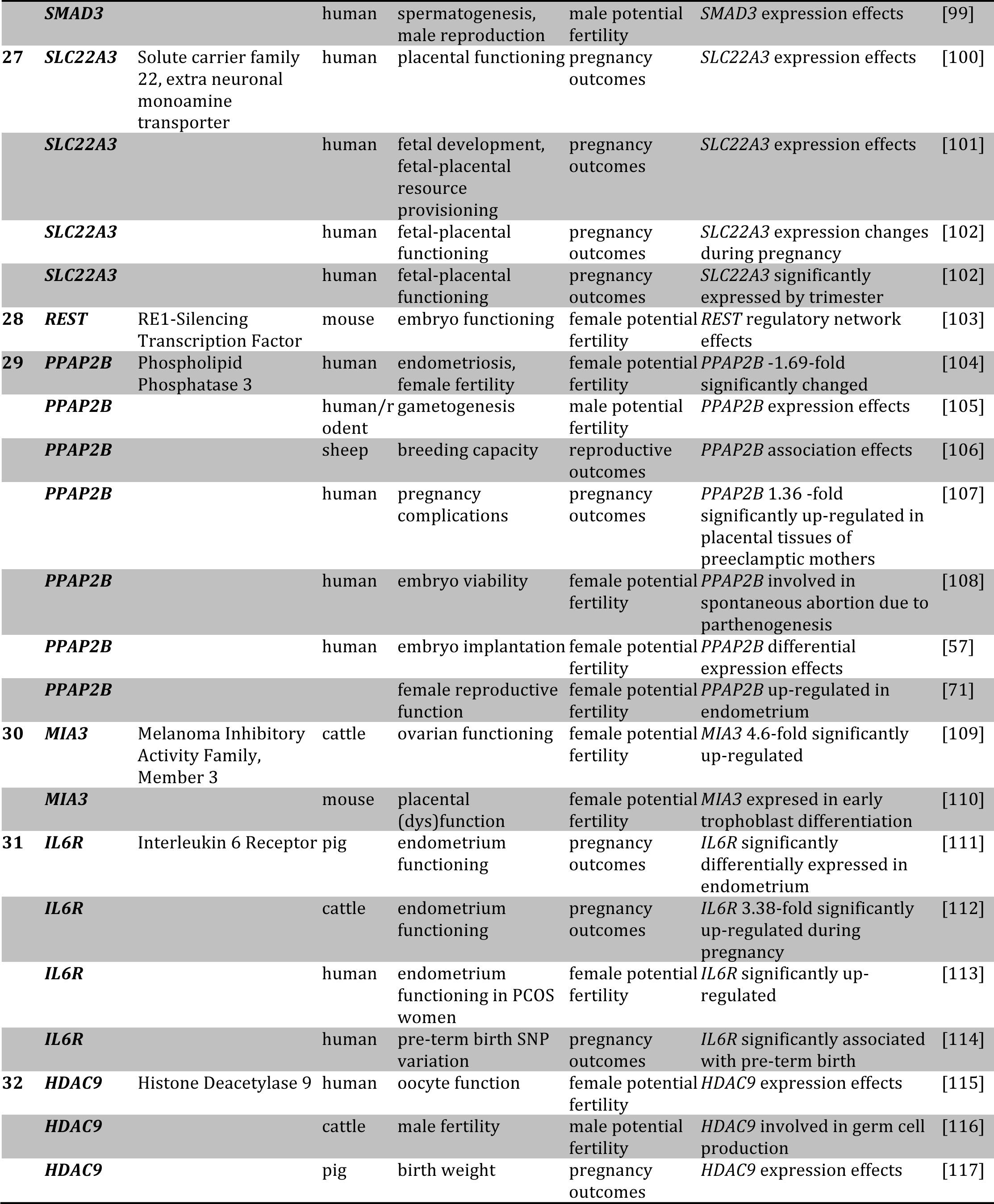
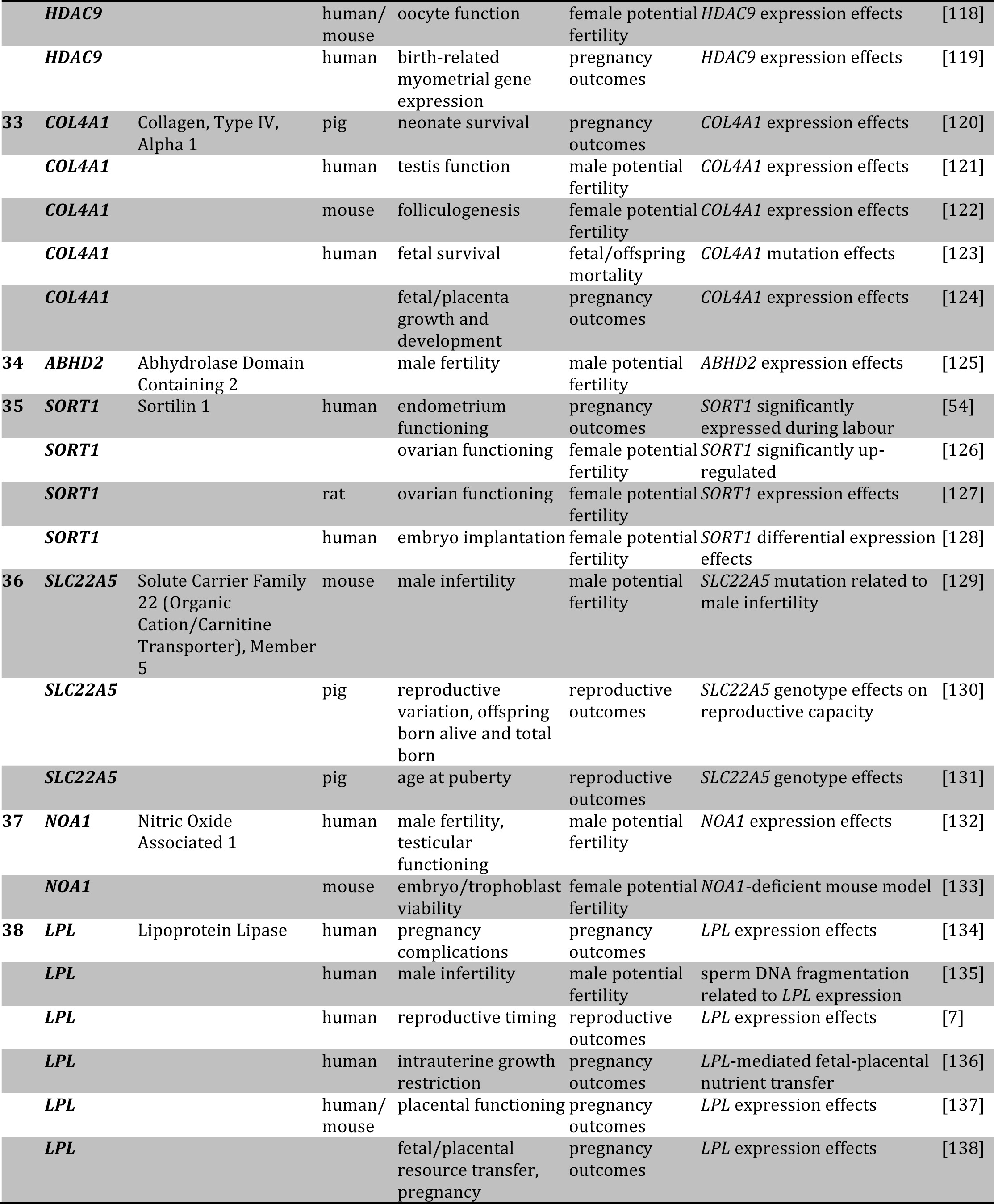
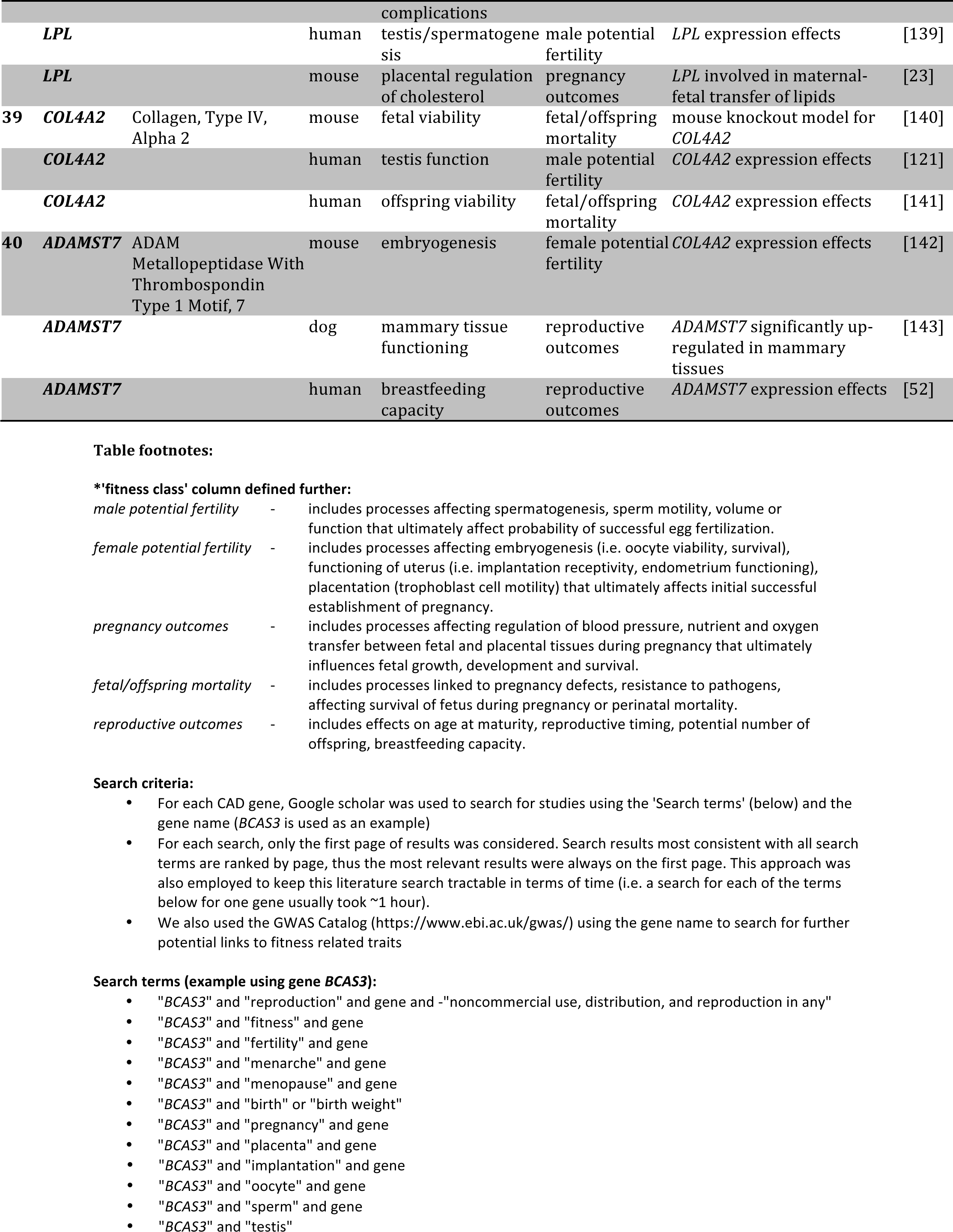
Pleiotropic links between coronary artery disease (CAD) and early-life fitness-related traits due to shared genetic loci. The table below provides extensive support (143 studies) that antagonistic pleiotropy is likely to be present for CAD genes due to their consistent connections with fitness-related traits expressed early in life. See Fig. 5 for discussion and conceptual overview of these potential effects. Fitness-related traits include fertility potential, reproductive outcomes, pregnancy outcomes, fetal growth and survival, i.e. affecting the ability of an organism to reproduce and transfer genes to the next generation. The first 3 columns give CAD gene rank (no.; based on rank of 40 genes from Fig. 1B), name and full name. Columns 4-8 provide key details of each study where CAD genes also contribute to traits that influence fitness, including what species that was demonstrated in, what biological process or fitness effects that gene is impacting, what fitness class that effect is likely to impact (e.g. dysfunctional spermatogenesis or embryogenesis will affect male and female fertility, ability to conceive), what the observed genetic effect or mechanism that gene was associated with.

**Table S2.**
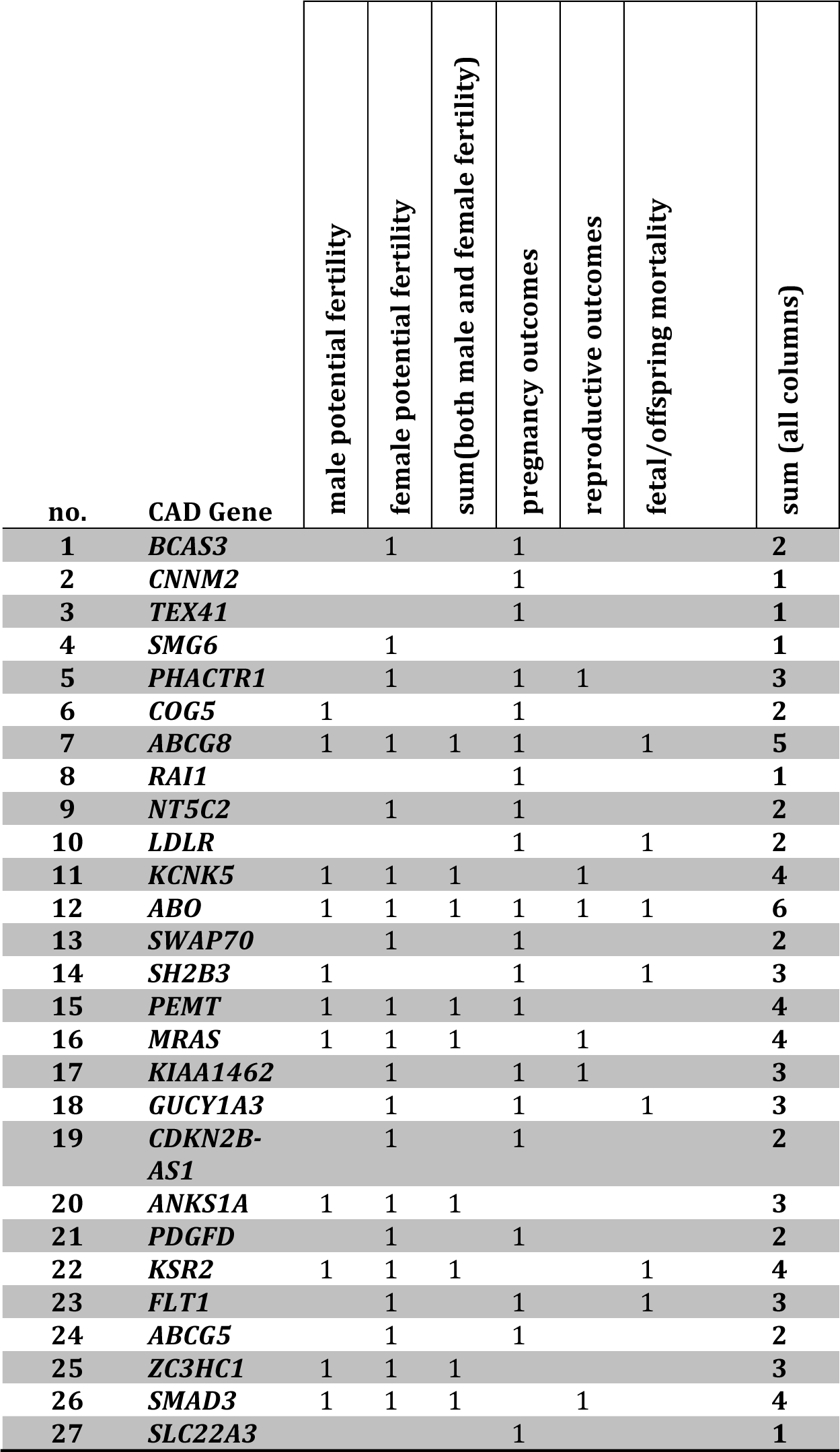
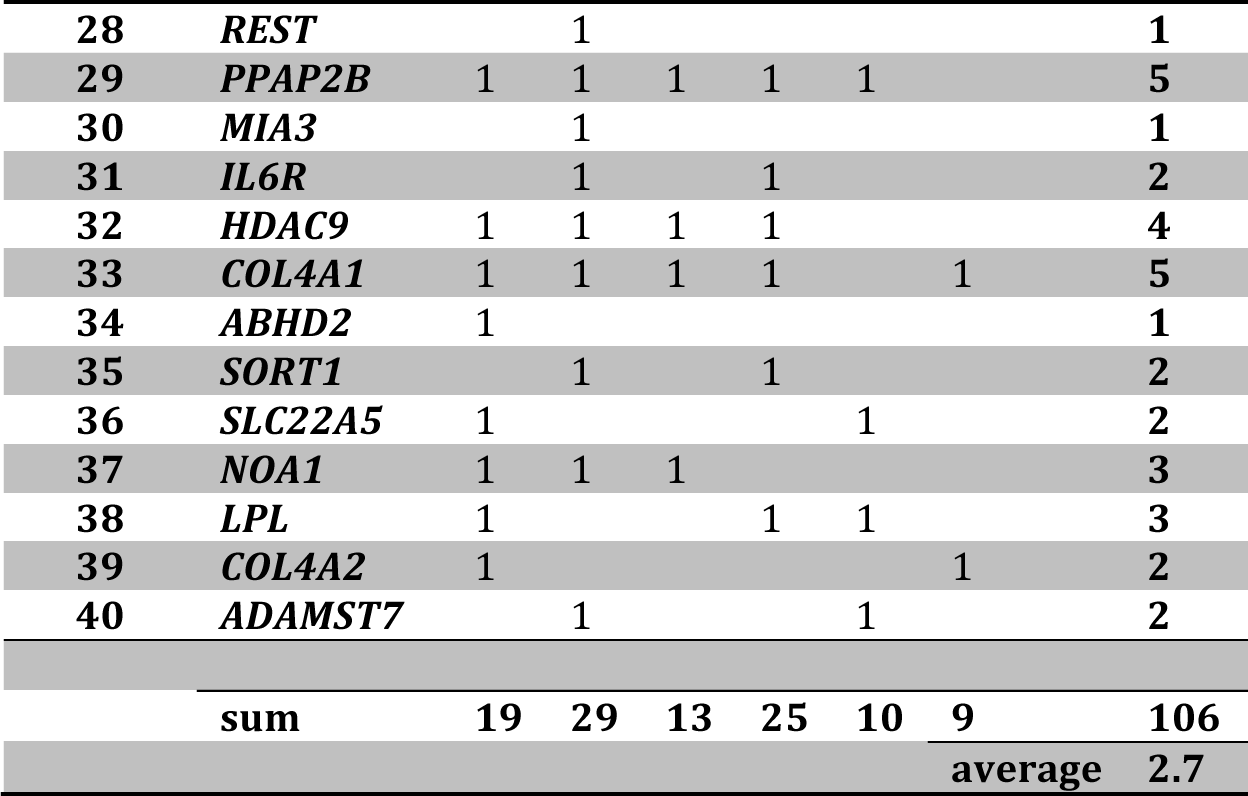
Summary of types of pleiotropic connections between coronary artery disease (CAD) and fitness-related traits. Counts are based on Table S1, ‘fitness class’ column. Most fitness-related traits were related to female potential fertility (29 of 40 genes had these effects) and pregnancy outcomes (25 of 40 genes had these effects). Some genes had broad or specific effects on fitness-related traits. For example, number of fitness classes affected ranged from 6 for *ABO* (had fitness effects across all classes) to 1, for example *CNNM2* (evidence for fitness effects in pregnancy outcomes class).

**Table S3.**
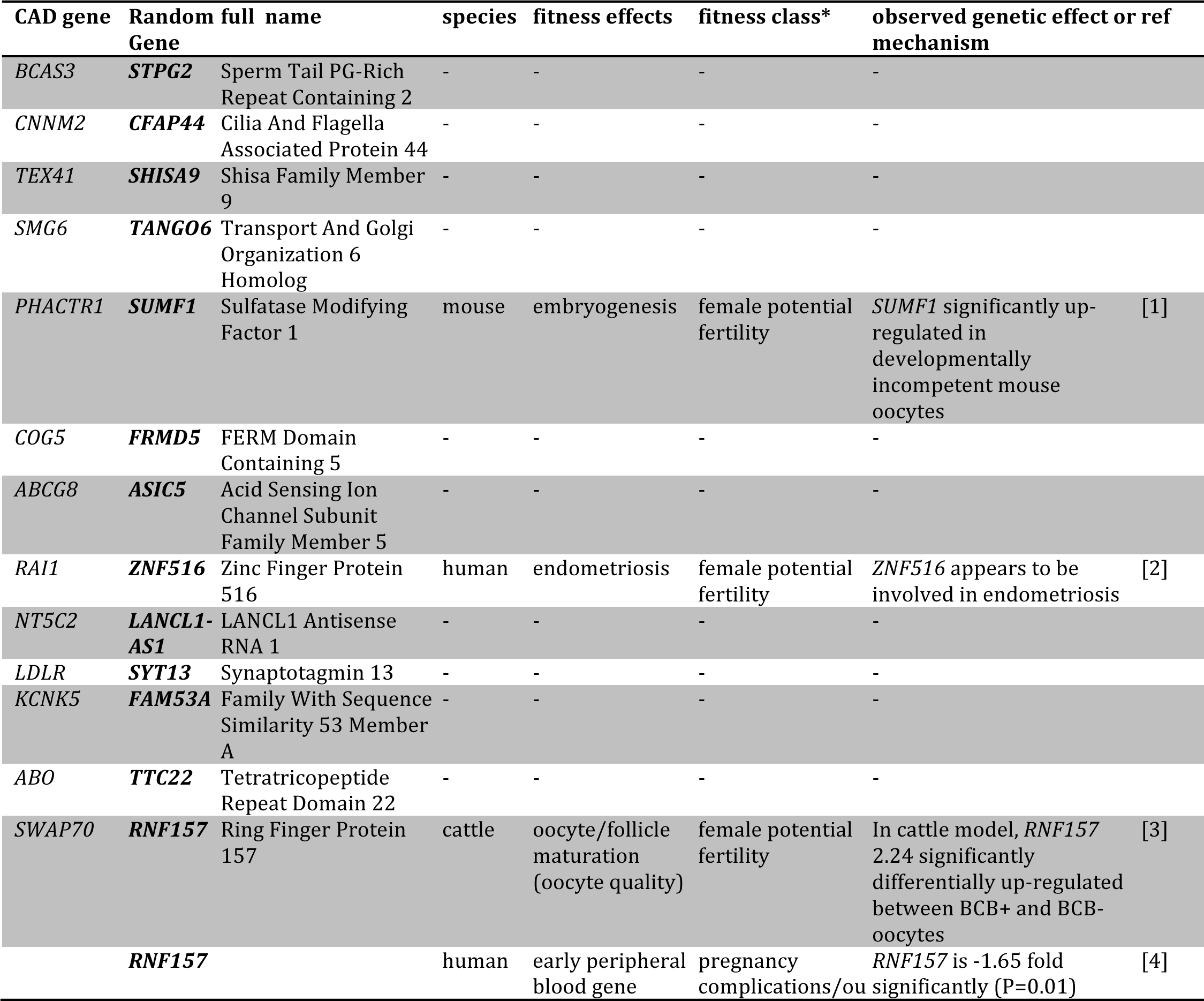
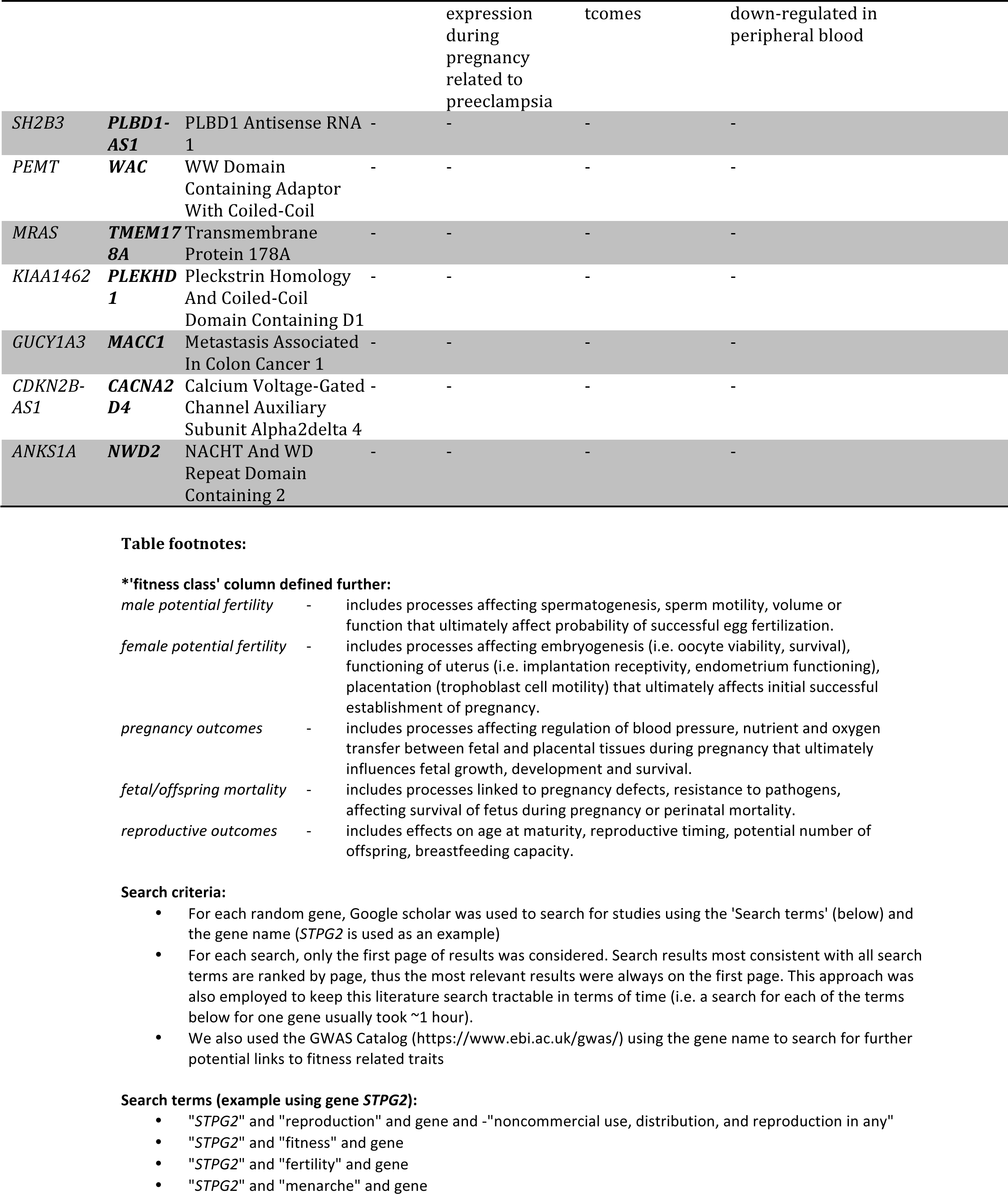
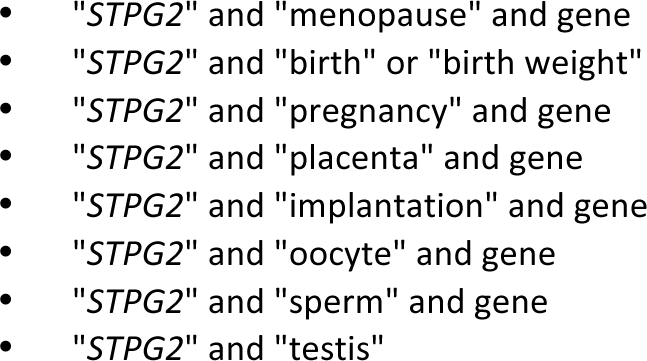
Pleiotropic links between randomly chosen genes and early-life fitness-related traits. Fitness-related traits include fertility potential, reproductive outcomes, pregnancy outcomes, fetal growth and survival, i.e. affecting the ability of an organism to reproduce and transfer genes to the next generation. The first column gives coronary artery disease (CAD) gene (first 20 of 40 CAD genes from Fig. 1B/Table S1). Columns 2-3 give name (abbreviated, full) of randomly chosen genes matched for approximate length for each CAD gene. Columns 4-8 provide key details of each study where random genes also contribute to traits that influence fitness, including what species that was demonstrated in, what biological process or fitness effects that gene is impacting, what fitness class that effect is likely to impact (e.g. dysfunctional spermatogenesis or embryogenesis will affect male and female fertility, ability to conceive), what the observed genetic effect or mechanism that gene was associated with.

**Table S4.**
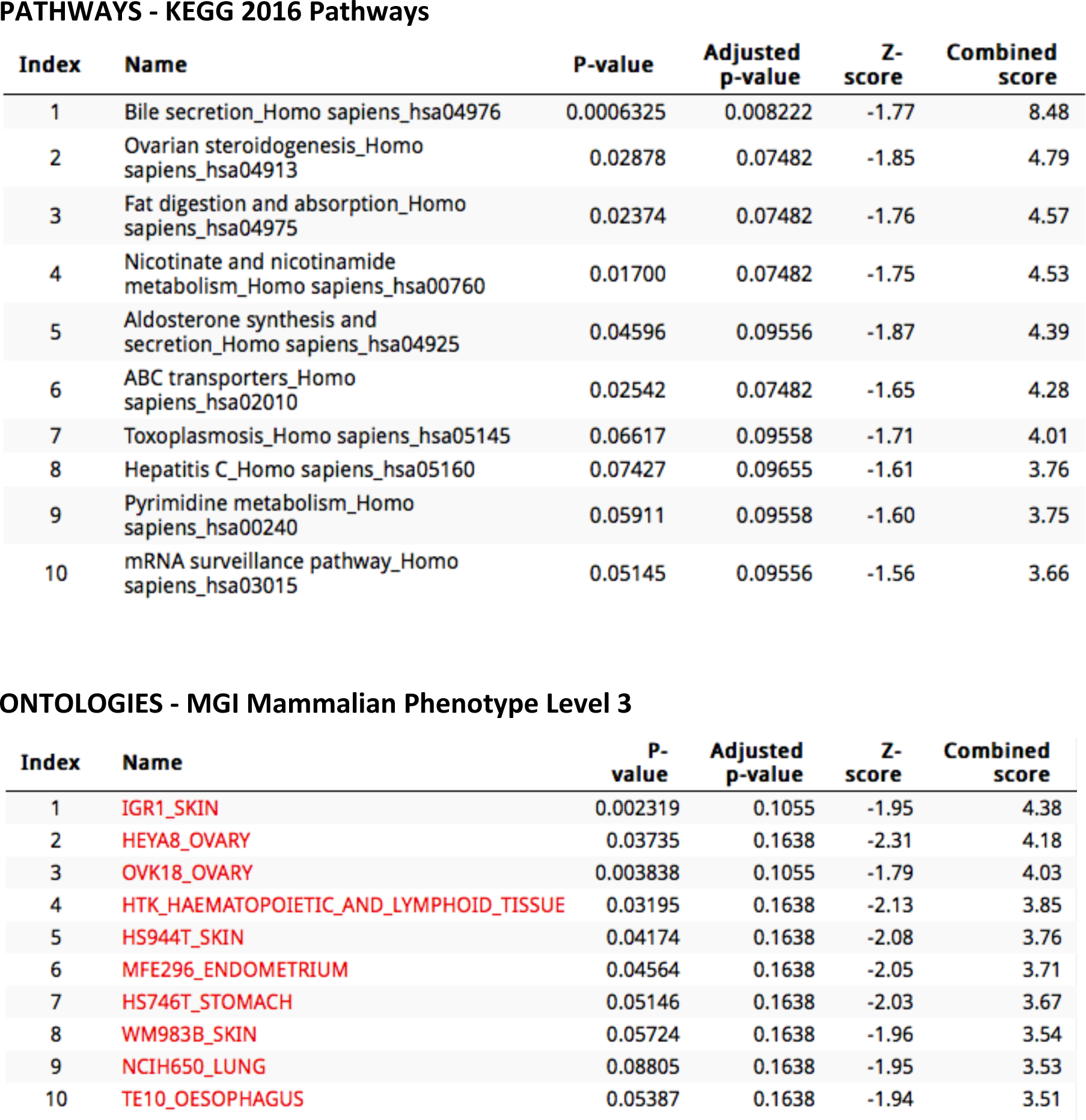
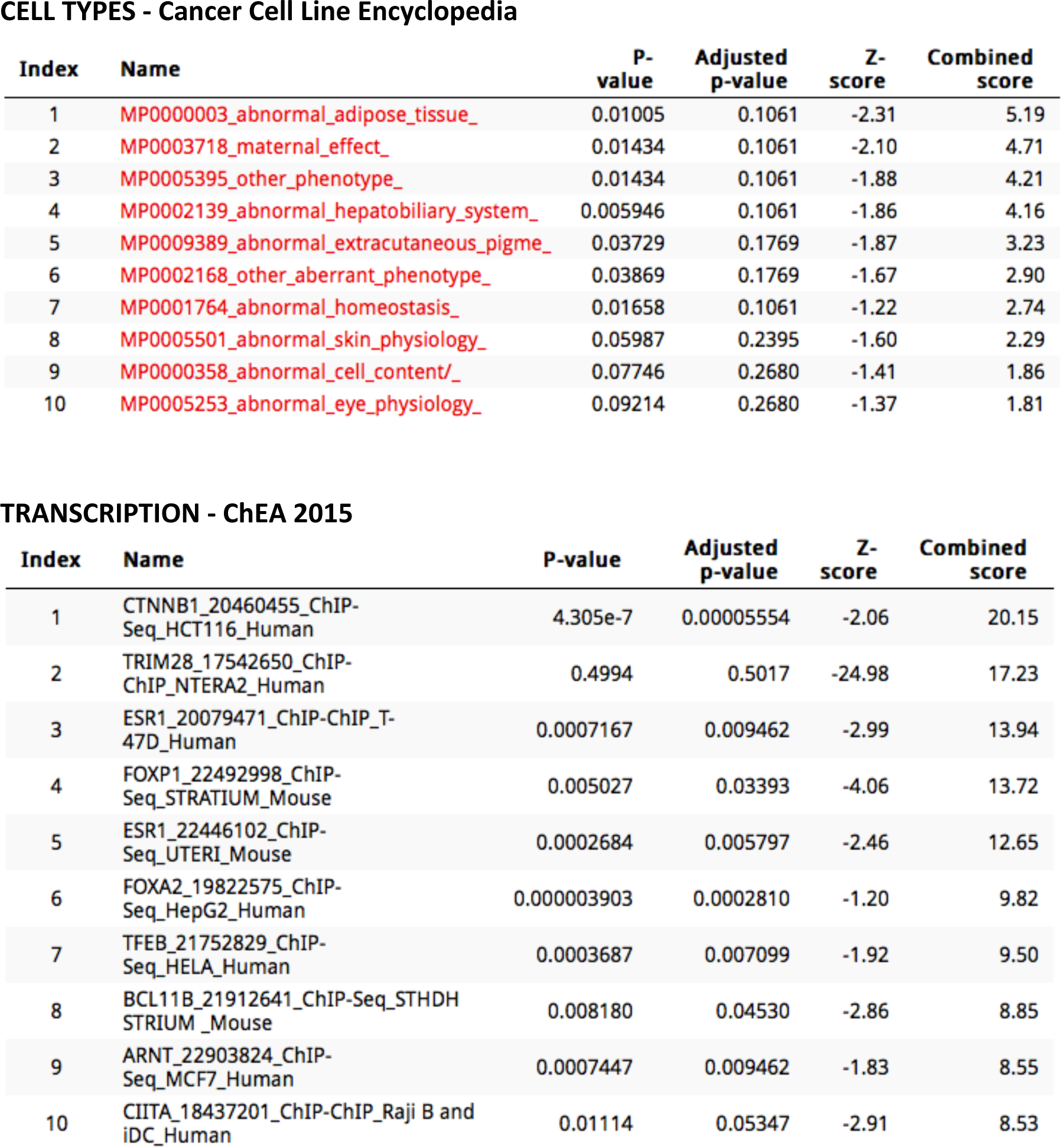
Selected Enrichr analysis outputs for top 10-ranked CAD genes with highest genetic risk-selection associations from Fig. 1B. Enrichr outputs includes KEGG 2016 Pathways (http://www.kegg.jp/kegg/download/), MGI Mammalian Phenotype Level 3 (http://www.informatics.jax.org/), Cancer Cell Line Encyclopaedia (http://portals.broadinstitute.org/ccle/data/browseData), and ChEA 2015 (http://amp.pharm.mssm.edu/lib/cheadownload.jsp).

## Supplementary Discussion

### Widespread signals of positive selection on CAD loci

Evidence of candidate positive selection signals for CAD loci were widespread, with many genes having significant iHS scores of small-medium size (i.e. iHS score range: 2-3) with four genes (*BCAS3*, *ANKS1A*, *CXCL12*, *PEMT*) harboring large selection signals (iHS >4), two of which had previously been identified as having strong selection signals including *BCAS3* (Breast Carcinoma Amplified Sequence 3) in the HapMap3 CEU population [1] and *PHACTR1* (phosphatase and actin regulator 1) across the ASW, CEU and CHB/CHD HapMap3 populations [2]. Twelve genes contained SNPs with selection scores that remained significant after correction for multiple testing (Fig. 1A). The consistency of smaller, less significant selection signals for several genes within most populations (i.e. *CNNM2*, *PHACTR1*, *PDGFD*) strongly suggest that these may be smaller and possibly valid incomplete selective sweeps that are typically missed due to stringency of multiple-correction thresholds and lack of validation across multiple populations.

These patterns match expectations from the polygenic model of selection that predicts that selection on complex traits mostly involves smaller shifts in many underlying loci; it is the likely reason why so few large selection signals have been found underlying complex traits in general [3, 4] and those underlying cardiovascular disease phenotypes in particular [5, 6]. For example, Kullo & Ding 2007 [6] found that 110 out of 364 genes in pathways associated with cardiovascular disease (i.e. inflammation, insulin, p53, Ras, cholesterol biosynthesis etc) had significantly higher Fst (empirical P<0.05) in at least one SNP between 4 populations, but none remained significant after correction for multiple testing. In a later study, Ding & Kullo 2011 [5] found that 8 out of 158 genome-wide significant SNPs in genes for 36 cardiovascular disease phenotypes and related traits (CHD, hypertension, stroke, BMI, lipids etc) had significantly elevated Fst between 52 populations in the Human Genome Diversity Project.

It is difficult to compare selection candidates we found in the 76 CAD associated genes with results from these two previous studies as full sets of gene lists and Fst estimates were not available for either, and they used loci underlying much broader cardiovascular disease phenotypes than our more current list of specific CAD loci [7]. Nevertheless, due to fine-scale imputation with the 1000 Genomes Panel, our study suggests that many more loci related to cardiovascular disease have been recently modified by natural selection than previously identified. The larger sample of SNPs also likely improved reliability of iHS p values, with many more estimates available per MAF bin used to standardize iHS measures [8].

The Fst measures used in the Ding and Kullo studies also differ qualitatively from the iHS scores we used. Fst captures allele frequency differences between populations and is less sensitive to detecting alleles that have undergone recent selection [9], while the iHS statistic detects whether common alleles are carried on unusually long haplotypes within populations and should be better at capturing more recent smaller selection signals [8]. Lastly, by considering not just genome-wide significant index SNPs, we were able to detect smaller selection signals within CAD loci that were consistent across populations and would have otherwise been missed. *PHACTR1* is a good example of this – several smaller candidate selection signals were found (iHS ranging from 2-3.8) where peak selection signals did not span the index SNP location - sometimes signals were in different introns within the same locus (Fig. S2).

## References

1. Akey, J.M., et al., Interrogating a high-density SNP map for signatures of natural selection. Genome Res, 2002. 12(12): p. 1805–14.

2. Bustamante, C.D., et al., Natural selection on protein-coding genes in the human genome. Nature, 2005. 437(7062): p. 1153–7.

3. Carlson, C.S., et al., Genomic regions exhibiting positive selection identified from dense genotype data. Genome Res, 2005. 15(11): p. 1553–65.

4. Kelley, J.L., et al., Genomic signatures of positive selection in humans and the limits of outlier approaches. Genome Res, 2006. 16(8): p. 980–9.

5. Lao, O., et al., Signatures of positive selection in genes associated with human skin pigmentation as revealed from analyses of single nucleotide polymorphisms. Annals of Human Genetics, 2007. 71: p. 354–369.

6. Sabeti, P.C., et al., Detecting recent positive selection in the human genome from haplotype structure. Nature, 2002. 419(6909): p. 832–7.

7. Shriver, M.D., et al., The genomic distribution of population substructure in four populations using 8,525 autosomal SNPs. Hum Genomics, 2004. 1(4): p. 274–86.

8. Tang, K., K.R. Thornton, and M. Stoneking, A new approach for using genome scans to detect recent positive selection in the human genome. Plos Biology, 2007. 5(7): p. 1587–1602.

9. Voight, B.F., et al., A map of recent positive selection in the human genome. PLoS Biol, 2006. 4(3): p. e72.

10. Grossman, S.R., et al., Identifying Recent Adaptations in Large-Scale Genomic Data. Cell, 2013. 152(4): p. 703–713.

11. Haasl, R.J. and B.A. Payseur, Fifteen years of genomewide scans for selection: trends, lessons and unaddressed genetic sources of complication. Mol Ecol, 2015.

12. Scheinfeldt, L.B. and S.A. Tishkoff, Recent human adaptation: genomic approaches, interpretation and insights. Nat Rev Genet, 2013. 14(10): p. 692–702.

13. Pritchard, J.K. and A. Di Rienzo, Adaptation - not by sweeps alone. Nature Reviews Genetics, 2010. 11(10): p. 665–667.

14. Pritchard, J.K., J.K. Pickrell, and G. Coop, The genetics of human adaptation: hard sweeps, soft sweeps, and polygenic adaptation. Curr Biol, 2010. 20(4): p. R208–15.

15. Kaplan, N.L., R.R. Hudson, and C.H. Langley, The Hitchhiking Effect Revisited. Genetics, 1989. 123(4): p. 887–899.

16. Smith, J.M. and J. Haigh, The hitch-hiking effect of a favourable gene. Genetics Research, 2007. 89(5-6): p. 391–403.

17. Oleksyk, T.K., M.W. Smith, and S.J. O’Brien, Genome-wide scans for footprints of natural selection. Philos Trans R Soc Lond B Biol Sci, 2010. 365(1537): p. 185–205.

18. Sabeti, P.C., et al., Positive natural selection in the human lineage. Science, 2006. 312(5780): p. 1614–20.

19. Wright, S., Genetical structure of populations. Nature, 1950. 166(4215): p. 247–9.

20. Hamblin, M.T., E.E. Thompson, and A. Di Rienzo, Complex signatures of natural selection at the Duffy blood group locus. American Journal of Human Genetics, 2002. 70(2): p. 369–383.

21. Beall, C.M., et al., Natural selection on EPAS1 (HIF2 alpha) associated with low hemoglobin concentration in Tibetan highlanders. Proceedings of the National Academy of Sciences of the United States of America, 2010. 107(25): p. 11459–11464.

22. Lamason, R.L., et al., SLC24A5, a putative cation exchanger, affects pigmentation in zebrafish and humans. Science, 2005. 310(5755): p. 1782–6.

23. Fraser, H.B., Gene expression drives local adaptation in humans. Genome Research, 2013. 23(7): p. 1089–1096.

24. Akey, J.M., Constructing genomic maps of positive selection in humans: where do we go from here? Genome Res, 2009. 19(5): p. 711–22.

25. Sabeti, P.C., et al., Genome-wide detection and characterization of positive selection in human populations. Nature, 2007. 449(7164): p. 913–U12.

26. Teshima, K.M., G. Coop, and M. Przeworski, How reliable are empirical genomic scans for selective sweeps? Genome Research, 2006. 16(6): p. 702–712.

27. Fu, W. and J.M. Akey, Selection and adaptation in the human genome. Annu Rev Genomics Hum Genet, 2013. 14: p. 467–89.

28. Hernandez, R.D., et al., Classic Selective Sweeps Were Rare in Recent Human Evolution. Science, 2011. 331(6019): p. 920–924.

29. Falconer, D.S. and T.F.C. Mackay, Introduction to quantitative genetics. 4th ed. 1996, Harlow, England; New York: Prentice Hall. xv, 464 p.

30. Grant, P.R. and B.R. Grant, Predicting Microevolutionary Responses to Directional Selection on Heritable Variation. Evolution, 1995. 49(2): p. 241-251.

31. Hermisson, J. and P.S. Pennings, Soft sweeps: Molecular population genetics of adaptation from standing genetic variation. Genetics, 2005. 169(4): p. 2335–2352.

32. Messer, P.W. and D.A. Petrov, Population genomics of rapid adaptation by soft selective sweeps. Trends in Ecology & Evolution, 2013. 28(11): p. 659–669.

33. Chevin, L.M. and F. Hospital, Selective Sweep at a Quantitative Trait Locus in the Presence of Background Genetic Variation. Genetics, 2008. 180(3): p. 1645–1660.

34. Welter, D., et al., The NHGRIGWAS Catalog, a curated resource of SNP-trait associations. Nucleic Acids Research, 2014. 42(D1): p. D1001–D1006.

35. Ding, K.Y. and I.J. Kullo, Geographic differences in allele frequencies of susceptibility SNPs for cardiovascular disease. Bmc Medical Genetics, 2011. 12.

36. Raj, T., et al., Common Risk Alleles for Inflammatory Diseases Are Targets of Recent Positive Selection. American Journal of Human Genetics, 2013. 92(4): p. 517–529.

37. Casto, A.M. and M.W. Feldman, Genome-Wide Association Study SNPs in the Human Genome Diversity Project Populations: Does Selection Affect Unlinked SNPs with Shared Trait Associations? Plos Genetics, 2011. 7(1).

38. Turchin, M.C., et al., Evidence of widespread selection on standing variation in Europe at height-associated SNPs. Nature Genetics, 2012. 44(9): p. 1015-+.

39. Go, A.S., et al., Heart disease and stroke statistics--2014 update: a report from the American Heart Association. Circulation, 2014. 129(3): p. e28–e292.

40. Deloukas, P., et al., Large-scale association analysis identifies new risk loci for coronary artery disease. Nature Genetics, 2013. 45(1): p. 25–U52.

41. Nikpay, M., et al., A comprehensive 1000 Genomes-based genome-wide association meta-analysis of coronary artery disease. Nature Genetics, 2015. 47(10): p. 1121-+.

42. Byars, S.G., et al., Colloquium papers: Natural selection in a contemporary human population. Proc Natl Acad Sci U S A, 2010. 107 Suppl 1: p. 1787–92.

43. Williams, G.C., Pleiotropy, Natural Selection, and the Evolution of Senescence. Evolution, 1957. 11(4): p. 398–411.

44. Jalowiec, D.A. and J.A. Hill, Myocardial infarction in the young and in women. Cardiovasc Clin, 1989. 20(1): p. 197–206.

45. Rubin, J.B. and W.B. Borden, Coronary Heart Disease in Young Adults. Current Atherosclerosis Reports, 2012. 14(2): p. 140–149.

46. Tuzcu, E.M., et al., High prevalence of coronary atherosclerosis in asymptomatic teenagers and young adults: evidence from intravascular ultrasound. Circulation, 2001. 103(22): p. 2705–10.

47. Morillas, P., et al., Characteristics and outcome of acute myocardial infarction in young patients - The PRIAMHO II study. Cardiology, 2007. 107(4): p. 217–225.

48. Allam, A.H., et al., Atherosclerosis in ancient Egyptian mummies: the Horus study. JACC Cardiovasc Imaging, 2011. 4(4): p. 315–27.

49. Wollstein, A. and W. Stephan, Inferring positive selection in humans from genomic data. Investig Genet, 2015. 6: p. 5.

50. Kuleshov, M.V., et al., Enrichr: a comprehensive gene set enrichment analysis web server 2016 update. Nucleic Acids Res, 2016. 44(W1): p. W90–7.

51. Kudaravalli, S., et al., Gene Expression Levels Are a Target of Recent Natural Selection in the Human Genome. Molecular Biology and Evolution, 2009. 26(3): p. 649–658.

52. Jarvis, J.P., et al., Patterns of Ancestry, Signatures of Natural Selection, and Genetic Association with Stature in Western African Pygmies. Plos Genetics, 2012. 8(4): p. 299–313.

53. Haritunians, T., et al., Genetic Predictors of Medically Refractory Ulcerative Colitis. Inflammatory Bowel Diseases, 2010. 16(11): p. 1830–1840.

54. Lang, B., et al., HDAC9 is implicated in schizophrenia and expressed specifically in post-mitotic neurons but not in adult neural stem cells. Am J Stem Cells, 2012. 1(1): p. 31–41.

55. Pokharel, K., et al., Transcriptome profiling of Finnsheep ovaries during out-of-season breeding period. Agricultural and Food Science, 2015. 24: p. 1–9.

56. Mbarek, H., et al., Identification of Common Genetic Variants Influencing Spontaneous Dizygotic Twinning and Female Fertility. Am J Hum Genet, 2016. 98(5): p. 898–908.

57. Huang, C., et al., Efficient SNP Discovery by Combining Microarray and Lab-on-a-Chip Data for Animal Breeding and Selection. Microarrays, 2015. 4(4): p. 570–595.

58. Mote, B.E., et al., Identification of genetic markers for productive life in commercial sows. J Anim Sci, 2009. 87(7): p. 2187–95.

59. Balgir, R.S., Menarcheal age in relation to ABO blood group phenotypes and haemoglobin-E genotypes. J Assoc Physicians India, 1993. 41(4): p. 210–1.

60. Pyun, J.A., et al., Genome-wide association studies and epistasis analyses of candidate genes related to age at menarche and age at natural menopause in a Korean population. Menopause, 2014. 21(5): p. 522–529.

61. Spencer, K.L., et al., Genetic Variation and Reproductive Timing: African American Women from the Population Architecture Using Genomics and Epidemiology (PAGE) Study. Plos One, 2013. 8(2).

62. Rempel, L.A., et al., Association analyses of candidate single nucleotide polymorphisms on reproductive traits in swine. J Anim Sci, 2010. 88(1): p. 1–15.

63. Patel, O.V., et al., Homeorhetic adaptation to lactation: comparative transcriptome analysis of mammary, liver, and adipose tissue during the transition from pregnancy to lactation in rats. Functional & Integrative Genomics, 2011. 11(1): p. 193–202.

64. Wang, M., et al., MicroRNA expression patterns in the bovine mammary gland are affected by stage of lactation. J Dairy Sci, 2012. 95(11): p. 6529–35.

65. Colodro-Conde, L., et al., A twin study of breastfeeding with a preliminary genome-wide association scan. Twin Res Hum Genet, 2015. 18(1): p. 61–72.

66. McLean, M.P., Z. Zhao, and G.C. Ness, Reduced hepatic LDL-receptor, 3-hydroxy-3-methylglutaryl coenzyme A reductase and sterol carrier protein-2 expression is associated with pregnancy loss in the diabetic rat. Endocrine, 1995. 3(10): p. 695–703.

67. Kuo, D.S., C. Labelle-Dumais, and D.B. Gould, COL4A1 and COL4A2 mutations and disease: insights into pathogenic mechanisms and potential therapeutic targets. Hum Mol Genet, 2012. 21(R1): p. R97–110.

68. Lin, S., et al., Pre-eclampsia has an adverse impact on maternal and fetal health. Transl Res, 2015. 165(4): p. 449–63.

69. Sayed, A.A.A., Molecular genetic studies in pregnancies affected by preeclampsia and intrauterine growth restriction. 2011, University of Nottingham.

70. Fritz, R.B., Trophoblast Retrieval and Isolation From e Cervix (tric) For NonInvasive Prenatal Genetic Diagnosis and Prediction Of Abnormal Pregnancy Outcome. 2015, Wayne State University.

71. Tabano, S., et al., Placental LPL gene expression is increased in severe intrauterine growth-restricted pregnancies. Pediatr Res, 2006. 59(2): p. 250–3.

72. Bhasin, K.K., et al., Maternal low-protein diet or hypercholesterolemia reduces circulating essential amino acids and leads to intrauterine growth restriction. Diabetes, 2009. 58(3): p. 559–66.

73. Kakourou, G., et al., Investigation of gene expression profiles before and after embryonic genome activation and assessment of functional pathways at the human metaphase II oocyte and blastocyst stage. Fertility and Sterility, 2013. 99(3): p. 803-+.

74. Siva, K., et al., Human BCAS3 Expression in Embryonic Stem Cells and Vascular Precursors Suggests a Role in Human Embryogenesis and Tumor Angiogenesis. Plos One, 2007. 2(11).

75. Liu, J., et al., Expression of SWAP-70 in the uterus and feto-maternal interface during embryonic implantation and pregnancy in the rhesus monkey (Macaca mulatta). Histochem Cell Biol, 2006. 126(6): p. 695–704.

76. Zhou, L., et al., Local injury to the endometrium in controlled ovarian hyperstimulation cycles improves implantation rates. Fertil Steril, 2008. 89(5): p. 1166–76.

77. Solca, C., G.S. Tint, and S.B. Patel, Dietary xenosterols lead to infertility and loss of abdominal adipose tissue in sterolin-deficient mice. J Lipid Res, 2013. 54(2): p. 397–409.

78. Moretti, E., et al., Ultrastructural study of spermatogenesis in KSR2 deficient mice. Transgenic Res, 2015. 24(4): p. 741–51.

79. Dokras, A., Cardiovascular disease risk in women with PCOS. Steroids, 2013. 78(8): p. 773–6.

80. Kenigsberg, S., et al., Gene expression microarray profiles of cumulus cells in lean and overweight-obese polycystic ovary syndrome patients. Mol Hum Reprod, 2009. 15(2): p. 89–103.

81. Manneras-Holm, L., A. Benrick, and E. Stener-Victorin, Gene expression in subcutaneous adipose tissue differs in women with polycystic ovary syndrome and controls matched pair-wise for age, body weight, and body mass index. Adipocyte, 2014. 3(3): p. 190–6.

82. Salilew-Wondim, D., et al., Polycystic ovarian syndrome is accompanied by repression of gene signatures associated with biosynthesis and metabolism of steroids, cholesterol and lipids. Journal of Ovarian Research, 2015. 8.

83. Scotti, L., et al., Platelet-derived growth factor BB and DD and angiopoietin1 are altered in follicular fluid from polycystic ovary syndrome patients. Mol Reprod Dev, 2014. 81(8): p. 748–56.

84. Yan, L., et al., Expression of apoptosis-related genes in the endometrium of polycystic ovary syndrome patients during the window of implantation. Gene, 2012. 506(2): p. 350–4.

85. Kilic, M., et al., Identification of Mutations and Evaluation of Cardiomyopathy in Turkish Patients with Primary Carnitine Deficiency. Jimd Reports - Case and Research Reports, 2011/3, 2012. 3: p. 17–23.

86. Tamai, I., Pharmacological and pathophysiological roles of carnitine/organic cation transporters (OCTNs: SLC22A4, SLC22A5 and Slc22a21). Biopharmaceutics & Drug Disposition, 2013. 34(1): p. 29–44.

87. Maqdasy, S., et al., Cholesterol and male fertility: What about orphans and adopted? Molecular and Cellular Endocrinology, 2013. 368(1-2): p. 30–46.

88. Schisterman, E.F., et al., Lipid concentrations and semen quality: the LIFE study. Andrology, 2014. 2(3): p. 408–15.

89. Carter, A.J. and A.Q. Nguyen, Antagonistic pleiotropy as a widespread mechanism for the maintenance of polymorphic disease alleles. BMC Med Genet, 2011. 12: p. 160.

90. Wang, X., S.G. Byars, and S.C. Stearns, Genetic links between post-reproductive lifespan and family size in Framingham. Evol Med Public Health, 2013. 2013(1): p. 241–53.

91. Granka, J.M., et al., Limited evidence for classic selective sweeps in African populations. Genetics, 2012. 192(3): p. 1049–64.

92. International HapMap, C., et al., A second generation human haplotype map of over 3.1 million SNPs. Nature, 2007. 449(7164): p. 851–61.

93. Surakka, I., et al., Founder population-specific HapMap panel increases power in GWA studies through improved imputation accuracy and CNV tagging. Genome Research, 2010. 20(10): p. 1344–1351.

94. Abraham, G. and M. Inouye, Fast principal component analysis of large-scale genome-wide data. PLoS One, 2014. 9(4): p. e93766.

95. Chang, C.C., et al., Second-generation PLINK: rising to the challenge of larger and richer datasets. Gigascience, 2015. 4: p. 7.

96. O’Connell, J., et al., A general approach for haplotype phasing across the full spectrum of relatedness. PLoS Genet, 2014. 10(4): p. e1004234.

97. Howie, B.N., P. Donnelly, and J. Marchini, A flexible and accurate genotype imputation method for the next generation of genome-wide association studies. PLoS Genet, 2009. 5(6): p. e1000529.

98. Paten, B., et al., Genome-wide nucleotide-level mammalian ancestor reconstruction. Genome Res, 2008. 18(11): p. 1829–43.

99. Flicek, P., et al., Ensembl 2012. Nucleic Acids Res, 2012. 40(Database issue): p. D84–90.

100. Gautier, M. and R. Vitalis, rehh: an R package to detect footprints of selection in genome-wide SNP data from haplotype structure. Bioinformatics, 2012. 28(8): p. 1176–7.

101. Stranger, B.E., et al., Patterns of Cis Regulatory Variation in Diverse Human Populations. Plos Genetics, 2012. 8(4): p. 272–284.

102. Stearns, S.C., The evolution of life histories. 1992, Oxford; New York: Oxford University Press. xii, 249 p.

103. Abraham, G., et al., Genomic prediction of coronary heart disease. bioRxiv, 2016.

## References

1. Siva, K., et al., human BCAS3 expression in embryonic stem cells and vascular precursors suggests a role in human embryogenesis and tumor angiogenesis. PLoS One, 2007. 2(11): p. e1202.

2. Zuccotti, M., et al., Maternal Oct-4 is a potential key regulator of the developmental competence of mouse oocytes. BMC Developmental Biology, 2008. 8(1): p. 1–14.

3. Nestler, A., et al., Blood pressure in pregnancy and magnesium sensitive genes. Pregnancy Hypertens, 2014. 4(1): p. 41–5.

4. Gheorghe, C.P., et al., Gene expression patterns in the hypoxic murine placenta: a role in epigenesis? Reprod Sci, 2007. 14(3): p. 223–33.

5. Yuan, H., et al., A de novo triplication on 2q22.3 including the entire ZEB2 gene associated with global developmental delay, multiple congenital anomalies and behavioral abnormalities. Mol Cytogenet, 2015. 8: p. 99.

6. Bao, J.Q., et al., UPF2, a nonsense-mediated mRNA decay factor, is required for prepubertal Sertoli cell development and male fertility by ensuring fidelity of the transcriptome. Development, 2015. 142(2): p. 352–362.

7. Spencer, K.L., et al., Genetic Variation and Reproductive Timing: African American Women from the Population Architecture Using Genomics and Epidemiology (PAGE) Study. Plos One, 2013. 8(2).

8. Kakourou, G., et al., Investigation of gene expression profiles before and after embryonic genome activation and assessment of functional pathways at the human metaphase II oocyte and blastocyst stage. Fertility and Sterility, 2013. 99(3): p. 803-+.

9. Saben, J., et al., Early growth response protein-1 mediates lipotoxicity-associated placental inflammation: role in maternal obesity. American Journal of Physiology-Endocrinology and Metabolism, 2013. 305(1): p. E1–E14.

10. Zhou, L., et al., Local injury to the endometrium in controlled ovarian hyperstimulation cycles improves implantation rates. Fertility and Sterility, 2008. 89(5): p. 1166–1176.

11. Newbold, R.R., et al., Developmental exposure to diethylstilbestrol alters uterine gene expression that may be associated with uterine neoplasia later in life. Molecular Carcinogenesis, 2007. 46(9): p. 783–796.

12. Patel, O.V., et al., Homeorhetic adaptation to lactation: comparative transcriptome analysis of mammary, liver, and adipose tissue during the transition from pregnancy to lactation in rats. Functional & Integrative Genomics, 2011. 11(1): p. 193–202.

13. Farkas, R.M., et al., The Drosophila Cog5 homologue is required for cytokinesis, cell elongation, and assembly of specialized golgi architecture during spermatogenesis. Molecular Biology of the Cell, 2003. 14(1): p. 190–200.

14. Foulquier, F., COG defects, birth and rise! Biochimica Et Biophysica Acta-Molecular Basis of Disease, 2009. 1792(9): p. 896–902.

15. Solca, C., G.S. Tint, and S.B. Patel, Dietary xenosterols lead to infertility and loss of abdominal adipose tissue in sterolin-deficient mice. Journal of Lipid Research, 2013. 54(2): p. 397–409.

16. Dixon, P.H. and C. Williamson, The pathophysiology of intrahepatic cholestasis of pregnancy. Clinics and Research in Hepatology and Gastroenterology, 2016. 40(2): p. 141–153.

17. Bi, W.M., et al., Inactivation of Rai1 in mice recapitulates phenotypes observed in chromosome engineered mouse models for Smith-Magenis syndrome. human Molecular Genetics, 2005. 14(8): p. 983–995.

18. Girirajan, S. and S.H. Elsea, Abnormal maternal behavior, altered sociability, and impaired serotonin metabolism in Rai1-transgenic mice. Mammalian Genome, 2009. 20(4): p. 247–255.

19. Shen, Z., et al., Estradiol regulation of nucleotidases in female reproductive tract epithelial cells and fibroblasts. PLoS One, 2013. 8(7): p. e69854.

20. Horikoshi, M., et al., New loci associated with birth weight identify genetic links between intrauterine growth and adult height and metabolism. Nat Genet, 2013. 45(1): p. 76–82.

21. Bhasin, K.K., et al., Maternal low-protein diet or hypercholesterolemia reduces circulating essential amino acids and leads to intrauterine growth restriction. Diabetes, 2009. 58(3): p. 559–66.

22. Ethier-Chiasson, M., et al., Influence of maternal lipid profile on placental protein expression of LDLr and SR-BI. Biochem Biophys Res Commun, 2007. 359(1): p. 8–14.

23. Overbergh, L., et al., Expression of mouse alpha-macroglobulins, lipoprotein receptor-related protein, LDL receptor, apolipoprotein E, and lipoprotein lipase in pregnancy. J Lipid Res, 1995. 36(8): p. 1774–86.

24. McLean, M.P., Z. Zhao, and G.C. Ness, Reduced hepatic LDL-receptor, 3-hydroxy-3-methylglutaryl coenzyme A reductase and sterol carrier protein-2 expression is associated with pregnancy loss in the diabetic rat. Endocrine, 1995. 3(10): p. 695–703.

25. Yeung, C.H. and T.G. Cooper, Potassium channels involved in human sperm volume regulation--quantitative studies at the protein and mRNA levels. Mol Reprod Dev, 2008. 75(4): p. 659–68.

26. Cooper, T.G. and C.H. Yeung, Involvement of potassium and chloride channels and other transporters in volume regulation by spermatozoa. Curr Pharm Des, 2007. 13(31): p. 3222–30.

27. Barfield, J.P., C.H. Yeung, and T.G. Cooper, Characterization of potassium channels involved in volume regulation of human spermatozoa. Mol Hum Reprod, 2005. 11(12): p. 891–7.

28. Chow, G.E., et al., Expression of two-pore domain potassium channels in nonhuman primate sperm. Fertil Steril, 2007. 87(2): p. 397–404.

29. Kang, D., et al., TASK-2 Expression Levels are Increased in mouse Cryopreserved Ovaries. Journal of Embryo Transfer, 2015. 30(4): p. 277–282.

30. Wang, M., et al., MicroRNA expression patterns in the bovine mammary gland are affected by stage of lactation. J Dairy Sci, 2012. 95(11): p. 6529–35.

31. Gloria-Bottini, F., et al., Effect of smoking and ABO blood groups on maternal age at child bearing and on birth weight. European Journal of Obstetrics & Gynecology and Reproductive Biology, 2011. 159(1): p. 83–86.

32. Clark, P. and I.A. Greer, The influence of maternal Lewis, Secretor and ABO(H) blood groups on fetal growth restriction. J Thromb Haemost, 2011. 9(12): p. 2411–5.

33. Dean, L., Hemolytic disease of the newborn, in Hemolytic disease of the newborn, L. Dean, Editor. 2005, National Center for Biotechnology Information (NCBI), National Library of Medicine, National Institutes of Health: Bethesda, MD.

34. Pathirana, S.L., et al., ABO-blood-group types and protection against severe, Plasmodium falciparum malaria. Ann Trop Med Parasitol, 2005. 99(2): p. 119–24.

35. Balgir, R.S., Menarcheal age in relation to ABO blood group phenotypes and haemoglobin-E genotypes. J Assoc Physicians India, 1993. 41(4): p. 210–1.

36. Omu, A.E., M. Al-Mutawa, and F. Al-Qattan, ABO blood group and expression of antisperm antibodies in infertile couples in Kuwait. Gynecol Obstet Invest, 1998. 45(1): p. 49–53.

37. Franchini, M., C. Mengoli, and G. Lippi, Relationship between ABO blood group and pregnancy complications: a systematic literature analysis. Blood Transfusion, 2016. 5: p. 1–8.

38. Mengoli, C., et al., ABO blood group and fertility: a single-centre study. Blood Transfus, 2015. 13(3): p. 521–3.

39. Liu, J., et al., Expression of SWAP-70 in the uterus and feto-maternal interface during embryonic implantation and pregnancy in the rhesus monkey (Macaca mulatta). Histochem Cell Biol, 2006. 126(6): p. 695–704.

40. Sitras, V., C. Fenton, and G. Acharya, Gene expression profile in cardiovascular disease and preeclampsia: a meta-analysis of the transcriptome based on raw data from human studies deposited in Gene Expression Omnibus. Placenta, 2015. 36(2): p. 170–8.

41. Padidela, R., Effects of polymorphisms in the growth hormone and insulin-like growth factor axis on intrauterine and postnatal growth. 2013, University College London.

42. Slack, C., et al., Regulation of lifespan, metabolism, and stress responses by the Drosophila SH2B protein, Lnk. PLoS Genet, 2010. 6(3): p. e1000881.

43. Hogeveen, M., et al., Umbilical choline and related methylamines betaine and dimethylglycine in relation to birth weight. Pediatr Res, 2013. 73(6): p. 783–7.

44. Zhu, J., Choline Intake During Pregnancy and Genetic Polymorphisms Influence Choline Metabolism in Chinese Preterms Receiving Total Parenteral Nutrition Therapy The Faseb Journal, 2016. 30.

45. Lazaros, L., et al., Phosphatidylethanolamine N-methyltransferase and choline dehydrogenase gene polymorphisms are associated with human sperm concentration. Asian J Androl, 2012. 14(5): p. 778–83.

46. Khot, V., et al., Expression of genes encoding enzymes involved in the one carbon cycle in rat placenta is determined by maternal micronutrients (folic acid, vitamin B12) and omega-3 fatty acids. Biomed Res Int, 2014. 2014: p. 613078.

47. Cheng, E.H., et al., Requirement of Leukemia Inhibitory Factor or Epidermal Growth Factor for Pre-Implantation Embryogenesis via JAK/STAT3 Signaling Pathways. PLoS One, 2016. 11(4): p. e0153086.

48. Yan, J., et al., Pregnancy alters choline dynamics: results of a randomized trial using stable isotope methodology in pregnant and nonpregnant women. Am J Clin Nutr, 2013. 98(6): p. 1459–67.

49. Gong, W., et al., Transcriptome profiling of the developing postnatal mouse testis using next-generation sequencing. Sci China Life Sci, 2013. 56(1): p. 1-12.

50. Cloke, B., et al., The androgen and progesterone receptors regulate distinct gene networks and cellular functions in decidualizing endometrium. Endocrinology, 2008. 149(9): p. 4462–74.

51. Palmqvist, L., et al., Correlation of murine embryonic stem cell gene expression profiles with functional measures of pluripotency. Stem Cells, 2005. 23(5): p. 663–80.

52. Colodro-Conde, L., et al., A twin study of breastfeeding with a preliminary genome-wide association scan. Twin Res Hum Genet, 2015. 18(1): p. 61–72.

53. Huang, C.-W., et al., Efficient SNP Discovery by Combining Microarray and Lab-on-a-Chip Data for Animal Breeding and Selection. Microarrays, 2015. 4(4): p. 570–595.

54. Chan, Y.W., et al., Assessment of myometrial transcriptome changes associated with spontaneous human labour by high-throughput RNA-seq. Exp Physiol, 2014. 99(3): p. 510–24.

55. Fledel-Alon, A., et al., Variation in human recombination rates and its genetic determinants. PLoS One, 2011. 6(6): p. e20321.

56. Perez-Montarelo, D., et al., Identification of genes regulating growth and fatness traits in pig through hypothalamic transcriptome analysis. Physiological Genomics, 2014. 46(6): p. 195–206.

57. Aghajanova, L., et al., Comparative Transcriptome Analysis of human Trophectoderm and Embryonic Stem Cell-Derived Trophoblasts Reveal Key Participants in Early Implantation. Biology of Reproduction, 2012. 86(1).

58. Ponsuksili, S., et al., Gene Expression and DNA-Methylation of Bovine Pretransfer Endometrium Depending on Its Receptivity after In Vitro-Produced Embryo Transfer. Plos One, 2012. 7(8).

59. Duzyj, C.M., et al., PreImplantation factor (PIF*) promotes embryotrophic and neuroprotective decidual genes: effect negated by epidermal growth factor. Journal of Neurodevelopmental Disorders, 2014. 6.

60. Sedlmeier, E.M., et al., human placental transcriptome shows sexually dimorphic gene expression and responsiveness to maternal dietary n-3 long-chain polyunsaturated fatty acid intervention during pregnancy. Bmc Genomics, 2014.15.

61. Taal, H.R., Early Growth, Cardiovascular and Renal Development - The Generation R Study. 2013, Erasmus University: Rotterdam.

62. Horikoshi, M., et al., New loci associated with birth weight identify genetic links between intrauterine growth and adult height and metabolism. Nature Genetics, 2013. 45(1): p. 76–U115.

63. Pagliardini, L., et al., Replication and meta-analysis of previous genome-wide association studies confirm vezatin as the locus with the strongest evidence for association with endometriosis. human Reproduction, 2015. 30(4): p. 987–993.

64. Sayed, A.-R.A.A.-R., Molecular genetic studies in pregnancies affected by preeclampsia and intrauterine growth restriction, in School of Molecular Medical Sciences. 2011, University of Nottingham: United Kingdom.

65. Moore, S.G., et al., Differentially Expressed Genes in Endometrium and Corpus Luteum of Holstein Cows Selected for High and Low Fertility Are Enriched for Sequence Variants Associated with Fertility. Biol Reprod, 2016. 94(1): p. 19.

66. Gad, A.Y.M.H., Transcriptome profiling of bovine blastocysts developed under alternative culture conditions during specific stages of development. 2012, University of Bonn: Bonn, Germany.

67. Tuttelmann, F., et al., Copy number variants in patients with severe oligozoospermia and Sertoli-cell-only syndrome. PLoS One, 2011. 6(4): p. e19426.

68. Scotti, L., et al., Platelet-derived growth factor BB and DD and angiopoietin1 are altered in follicular fluid from polycystic ovary syndrome patients. Mol Reprod Dev, 2014. 81(8): p. 748–56.

69. Bonnet, A., et al., Spatio-Temporal Gene Expression Profiling during In Vivo Early Ovarian Folliculogenesis: Integrated Transcriptomic Study and Molecular Signature of Early Follicular Growth. PLoS One, 2015. 10(11): p. e0141482.

70. Sharov, A.A., et al., Effects of aging and calorie restriction on the global gene expression profiles of mouse testis and ovary. BMC Biol, 2008. 6: p. 24.

71. Popovici, R.M., et al., Gene expression profiling of human endometrial-trophoblast interaction in a coculture model. Endocrinology, 2006. 147(12): p. 5662–5675.

72. Sleer, L.S. and C.C. Taylor, Cell-type localization of platelet-derived growth factors and receptors in the postnatal rat ovary and follicle. Biol Reprod, 2007. 76(3): p. 379–90.

73. Sitras, V., et al., Differential placental gene expression in severe preeclampsia. Placenta, 2009. 30(5): p. 424–33.

74. Moretti, E., et al., Ultrastructural study of spermatogenesis in KSR2 deficient mice. Transgenic Res, 2015. 24(4): p. 741–51.

75. Cerny, K.L., et al., A transcriptomal analysis of bovine oviductal epithelial cells collected during the follicular phase versus the luteal phase of the estrous cycle. Reprod Biol Endocrinol, 2015. 13: p. 84.

76. Henry, M.D., et al., Obesity-dependent dysregulation of glucose homeostasis in kinase suppressor of ras 2-/-mice. Physiol Rep, 2014. 2(7).

77. Kaipainen, A., et al., The Related Flt4, Flt1, and Kdr Receptor Tyrosine Kinases Show Distinct Expression Patterns in human Fetal Endothelial-Cells. Journal of Experimental Medicine, 1993. 178(6): p. 2077–2088.

78. Khankin, E.V., et al., Hemodynamic, Vascular, and Reproductive Impact of FMS-Like Tyrosine Kinase 1 (FLT1) Blockade on the Uteroplacental Circulation During Normal mouse Pregnancy. Biology of Reproduction, 2012. 86(2).

79. Muehlenbachs, A., et al., Natural selection of FLT1 alleles and their association with malaria resistance in utero. Proceedings of the National Academy of Sciences of the United States of America, 2008. 105(38): p. 14488–14491.

80. Bonnet, A., et al., An overview of gene expression dynamics during early ovarian folliculogenesis: specificity of follicular compartments and bidirectional dialog. Bmc Genomics, 2013. 14.

81. Fritz, R., Trophoblast Retrieval And Isolation From The Cervix (tric) For NonInvasive Prenatal Genetic Diagnosis And Prediction Of Abnormal Pregnancy Outcome, in Physiology. 2015, Wayne State University.

82. Korevaar, T.I., et al., Soluble Flt1 and placental growth factor are novel determinants of newborn thyroid (dys)function: the generation R study. J Clin Endocrinol Metab, 2014. 99(9): p. E1627–34.

83. Dan, S., L. Haibo, and L. Hong, Pathogenesis and stem cell therapy for premature ovarian failure. OA Stem Cells, 2014. 10(2).

84. Dey, S.K., et al., Molecular cues to implantation. Endocr Rev, 2004. 25(3): p. 341–73.

85. Sood, R., et al., Gene expression patterns in human placenta. Proc Natl Acad Sci U S A, 2006. 103(14): p. 5478–83.

86. Tsao, P.N., et al., Excess soluble fms-like tyrosine kinase 1 and low platelet counts in premature neonates of preeclamptic mothers. Pediatrics, 2005. 116(2): p. 468–72.

87. Chen, L.H., et al., [Effects of intrauterine growth restriction and high-fat diet on serum lipid and transcriptional levels of related hepatic genes in rats]. Zhongguo Dang Dai Er Ke Za Zhi, 2015. 17(10): p. 1124–30.

88. Ozawa, M., et al., Global gene expression of the inner cell mass and trophectoderm of the bovine blastocyst. BMC Dev Biol, 2012. 12: p. 33.

89. Archambeault, D.R. and M.M. Matzuk, Disrupting the male germ line to find infertility and contraception targets. Ann Endocrinol (Paris), 2014. 75(2): p. 101–8.

90. Niklaus, A.L. and J.W. Pollard, Mining the mouse transcriptome of receptive endometrium reveals distinct molecular signatures for the luminal and glandular epithelium. Endocrinology, 2006. 147(7): p. 3375–90.

91. Xu, J., J. Oakley, and E.A. McGee, Stage-specific expression of Smad2 and Smad3 during folliculogenesis. Biol Reprod, 2002. 66(6): p. 1571–8.

92. Ethier, J.F. and J.K. Findlay, Roles of activin and its signal transduction mechanisms in reproductive tissues. Reproduction, 2001. 121(5): p. 667–75.

93. Matsuda, T., et al., Cross-talk between transforming growth factor-beta and estrogen receptor signaling through Smad3. Journal of Biological Chemistry, 2001. 276(46): p. 42908–42914.

94. Xu, R., et al., Developmental and stage-specific expression of Smad2 and Smad3 in rat testis. Journal of Andrology, 2003. 24(2): p. 192–200.

95. Pyun, J.A., et al., Genome-wide association studies and epistasis analyses of candidate genes related to age at menarche and age at natural menopause in a Korean population. Menopause-the Journal of the North American Menopause Society, 2014. 21(5): p. 522–529.

96. Mbarek, H., et al., Identification of Common Genetic Variants Influencing Spontaneous Dizygotic Twinning and Female Fertility. American Journal of human Genetics, 2016. 98(5): p. 898–908.

97. Liu, Y.X., et al., Effects of Smad3 on the Proliferation and Steroidogenesis in human Ovarian Luteinized Granulosa Cells. Iubmb Life, 2014. 66(6): p. 424–437.

98. Dunn, N.R., et al., Combinatorial activities of Smad2 and Smad3 regulate mesoderm formation and patterning in the mouse embryo. Development, 2004. 131(8): p. 1717–1728.

99. Itman, C., et al., Smad3 Dosage Determines Androgen Responsiveness and Sets the Pace of Postnatal Testis Development. Endocrinology, 2011. 152(5): p. 2076–2089.

100. Jacob, S. and K.H. Moley, Gametes and embryo epigenetic reprogramming affect developmental outcome: Implication for assisted reproductive technologies. Pediatric Research, 2005. 58(3): p. 437–446.

101. Nelissen, E.C.M., et al., Epigenetics and the placenta. human Reproduction Update, 2011. 17(3): p. 397–417.

102. Monk, D., et al., Limited evolutionary conservation of imprinting in the human placenta. Proceedings of the National Academy of Sciences of the United States of America, 2006. 103(17): p. 6623–6628.

103. Johnson, R., et al., REST Regulates Distinct Transcriptional Networks in Embryonic and Neural Stem Cells. Plos Biology, 2008. 6(10): p. 2205–2219.

104. Burney, R.O., et al., Gene expression analysis of endometrium reveals progesterone resistance and candidate susceptibility genes in women with endometriosis. Endocrinology, 2007. 148(8): p. 3814–3826.

105. Chalmel, F., et al., The conserved transcriptome in human and rodent male gametogenesis. Proceedings of the National Academy of Sciences of the United States of America, 2007. 104(20): p. 8346–8351.

106. Pokharel, K., et al., Transcriptome profiling of Finnsheep ovaries during out-of-season breeding period. Agricultural and Food Science, 2015. 24(1): p. 1–9.

107. Yan, Y.H., et al., Screening for preeclampsia pathogenesis related genes. European Review for Medical and Pharmacological Sciences, 2013. 17(22): p. 3083–3094.

108. Liu, N., et al., Genome-wide Gene Expression Profiling Reveals Aberrant MAPK and Wnt Signaling Pathways Associated with Early Parthenogenesis. Journal of Molecular Cell Biology, 2010. 2(6): p. 333–344.

109. Hatzirodos, N., et al., Transcriptome profiling of granulosa cells from bovine ovarian follicles during atresia. Bmc Genomics, 2014. 15.

110. Moore, A.C. and A. Sutherland, Characterizing the Nature of the Xst199 Mutation and the Normal Function of the TANGO1 Protein in Trophoblast Cells. Faseb Journal, 2011. 25.

111. Ostrup, E., et al., Differential Endometrial Gene Expression in Pregnant and Nonpregnant Sows. Biology of Reproduction, 2010. 83(2): p. 277–285.

112. Mansouri-Attia, N., et al., Endometrium as an early sensor of in vitro embryo manipulation technologies. Proceedings of the National Academy of Sciences of the United States of America, 2009. 106(14): p. 5687–5692.

113. Yan, L., et al., Expression of apoptosis-related genes in the endometrium of polycystic ovary syndrome patients during the window of implantation. Gene, 2012. 506(2): p. 350–354.

114. Anum, E.A., et al., Genetic Contributions to Disparities in Preterm Birth. Pediatric Research, 2009. 65(1): p. 1–9.

115. Zhang, P., et al., Distinct sets of developmentally regulated genes that are expressed by human oocytes and human embryonic stem cells. Fertility and Sterility, 2007. 87(3): p. 677–690.

116. Hering, D.M., K. Olenski, and S. Kaminski, Genome-Wide Association Study for Sperm Concentration in Holstein-Friesian Bulls. Reproduction in Domestic Animals, 2014. 49(6): p. 1008–1014.

117. Zhang, L.F., et al., Genome Wide Screening of Candidate Genes for Improving piglet Birth Weight Using High and Low Estimated Breeding Value Populations. International Journal of Biological Sciences, 2014. 10(3): p. 236–244.

118. Assou, S., et al., The human cumulus-oocyte complex gene-expression profile. human Reproduction, 2006. 21(7): p. 1705–1719.

119. Chan, Y.W., et al., Assessment of myometrial transcriptome changes associated with spontaneous human labour by high-throughput RNA-seq. Experimental Physiology, 2014. 99(3): p. 510–524.

120. Jiang, L., et al., Expression of X-linked genes in deceased Neonates and surviving cloned female piglets. Molecular Reproduction and Development, 2008. 75(2): p. 265–273.

121. Chen, S.R., et al., The Wilms Tumor Gene, Wt1, Maintains Testicular Cord Integrity by Regulating the Expression of Col4a1 and Col4a2. Biology of Reproduction, 2013. 88(3).

122. Borup, R., et al., Competence Classification of Cumulus and Granulosa Cell Transcriptome in Embryos Matched by Morphology and Female Age. Plos One, 2016. 11(4).

123. Garel, C., et al., Fetal intracerebral hemorrhage and COL4A1 mutation: promise and uncertainty. Ultrasound in Obstetrics & Gynecology, 2013. 41(2): p. 228–230.

124. Rodriguez-Zas, S.L., K. Schellander, and H.A. Lewin, Biological interpretations of transcriptomic profiles in mammalian oocytes and embryos. Reproduction, 2008. 135(2): p. 129–139.

125. Gerhardt, K., Progesterone and Endocannabinoid Interaction Alters Sperm Activation. Biol Reprod, 2016. 95(1): p. 9.

126. Zhao, Y., et al., Effect of luteal-phase support on endometrial microRNA expression following controlled ovarian stimulation. Reprod Biol Endocrinol, 2012. 10: p. 72.

127. Wang, C.Q. and L.-S. Ll, Effect of CdCl_2 on Expression of Sortilin(Sort1) in rat Ovary. Reproduction and Contraception, 2012. 7.

128. Aghajanova, L., et al., Comparative transcriptome analysis of human trophectoderm and embryonic stem cell-derived trophoblasts reveal key participants in early implantation. Biol Reprod, 2012. 86(1): p. 1–21.

129. Toshimori, K., et al., Dysfunctions of the epididymis as a result of primary carnitine deficiency in juvenile visceral steatosis mice. Febs Letters, 1999. 446(2-3): p. 323–326.

130. Mote, B.E., et al., Identification of genetic markers for productive life in commercial sows. Journal of Animal Science, 2009. 87(7): p. 2187–2195.

131. Rempel, L.A., et al., Association analyses of candidate single nucleotide polymorphisms on reproductive traits in swine. Journal of Animal Science, 2010. 88(1): p. 1–15.

132. Okada, H., et al., Genome-wide expression of azoospermia testes demonstrates a specific profile and implicates ART3 in genetic susceptibility. Plos Genetics, 2008. 4(2).

133. Kolanczyk, M., et al., NOA1 is an essential GTPase required for mitochondrial protein synthesis. Molecular Biology of the Cell, 2011. 22(1): p. 1–11.

134. Schmella, M.J., et al., The-93T/G LPL Promoter Polymorphism Is Associated With Lower Third-Trimester Triglycerides in Pregnant African American Women. Biological Research for Nursing, 2015. 17(4): p. 429–437.

135. Intasqui, P., et al., Sperm nuclear DNA fragmentation rate is associated with differential protein expression and enriched functions in human seminal plasma. Bju International, 2013. 112(6): p. 835–843.

136. Tabano, S., et al., Placental LPL gene expression is increased in severe intrauterine growth-restricted pregnancies. Pediatric Research, 2006. 59(2): p. 250–253.

137. Lindegaard, M.L.S., et al., Endothelial and lipoprotein lipases in human and mouse placenta. Journal of Lipid Research, 2005. 46(11): p. 2339–2346.

138. Lager, S. and T.L. Powell, Regulation of nutrient transport across the placenta. J Pregnancy, 2012. 2012: p. 179827.

139. Nielsen, J.E., et al., Lipoprotein lipase and endothelial lipase in human testis and in germ cell neoplasms. Int J Androl, 2010. 33(1): p. e207–15.

140. Kuo, D.S., C. Labelle-Dumais, and D.B. Gould, COL4A1 and COL4A2 mutations and disease: insights into pathogenic mechanisms and potential therapeutic targets. human Molecular Genetics, 2012. 21: p. R97–R110.

141. Yoneda, Y., et al., De novo and inherited mutations in COL4A2, encoding the type IV collagen alpha2 chain cause porencephaly. Am J Hum Genet, 2012. 90(1): p. 86–90.

142. Hurskainen, T.L., et al., ADAM-TS5, ADAM-TS6, and ADAM-TS7, novel members of a new family of zinc metalloproteases. General features and genomic distribution of the ADAM-TS family. J Biol Chem, 1999. 274(36): p. 25555–63.

143. Rao, N.A., et al., Gene expression profiles of progestin-induced canine mammary hyperplasia and spontaneous mammary tumors. J Physiol Pharmacol, 2009. 60 Suppl 1: p. 73–84.

## References

1. Zuccotti, M., et al., Maternal Oct-4 is a potential key regulator of the developmental competence of mouse oocytes. BMC Developmental Biology, 2008. 8(1): p. 1–14.

2. Sun, P.R., et al., Genome-wide profiling of long noncoding ribonucleic acid expression patterns in ovarian endometriosis by microarray. Fertil Steril, 2014. 101(4): p. 1038–46 e7.

3. Janowski, D., et al., Incidence of apoptosis and transcript abundance in bovine follicular cells is associated with the quality of the enclosed oocyte. Theriogenology, 2012. 78(3): p. 656–659.

4. Enquobahrie, D.A., et al., Maternal peripheral blood gene expression in early pregnancy and preeclampsia. Int J Mol Epidemiol Genet, 2011. 2(1): p. 78–94.

## References

1. Sabeti, P.C., et al., Genome-wide detection and characterization of positive selection in human populations. Nature, 2007. 449(7164): p. 913–U12.

2. Williamson, S.H., et al., Localizing recent adaptive evolution in the human genome. Plos Genetics, 2007. 3(6): p. 901–915.

3. Fu, W. and J.M. Akey, Selection and adaptation in the human genome. Annu Rev Genomics Hum Genet, 2013. 14: p. 467–89.

4. Hernandez, R.D., et al., Classic Selective Sweeps Were Rare in Recent Human Evolution. Science, 2011. 331(6019): p. 920–924.

5. Ding, K.Y. and I.J. Kullo, Geographic differences in allele frequencies of susceptibility SNPs for cardiovascular disease. Bmc Medical Genetics, 2011. 12.

6. Kullo, I.J. and K.Y. Ding, Patterns of population differentiation of candidate genes for cardiovascular disease. Bmc Genetics, 2007. 8.

7. Nikpay, M., et al., A comprehensive 1000 Genomes-based genome-wide association meta-analysis of coronary artery disease. Nature Genetics, 2015. 47(10): p. 1121-+.

8. Voight, B.F., et al., A map of recent positive selection in the human genome. PLoS Biol, 2006. 4(3): p. e72.

9. Sabeti, P.C., et al., Positive natural selection in the human lineage. Science, 2006. 312(5780): p. 1614–20.

